# Promoter sequence and architecture determine expression variability and confer robustness to genetic variants

**DOI:** 10.1101/2021.10.29.466407

**Authors:** Hjörleifur Einarsson, Marco Salvatore, Christian Vaagensø, Nicolas Alcaraz, Jette Bornholdt, Sarah Rennie, Robin Andersson

## Abstract

Genetic and environmental exposures cause variability in gene expression. Although most genes are affected in a population, their effect sizes vary greatly, indicating the existence of regulatory mechanisms that could amplify or attenuate expression variability. Here, we investigate the relationship between the sequence and transcription start site architectures of promoters and their expression variability across human individuals. We find that expression variability can be largely explained by a promoter’s DNA sequence and its binding sites for specific transcription factors. We show that promoter expression variability reflects the biological process of a gene, demonstrating a selective trade-off between stability for metabolic genes and plasticity for responsive genes and those involved in signaling. Promoters with a rigid transcription start site architecture are more prone to have variable expression and to be associated with genetic variants with large effect sizes, while a flexible usage of transcription start sites within a promoter attenuates expression variability and limits genotypic effects. Our work provides insights into the variable nature of responsive genes and reveals a novel mechanism for supplying transcriptional and mutational robustness to essential genes through multiple transcription start site regions within a promoter.

## Introduction

Transcriptional regulation is the main process controlling how genome-encoded information is translated into phenotypes. Hence, understanding how transcriptional regulation influences gene expression variability is of fundamental importance to understand how organisms are capable of generating proper phenotypes in the face of stochastic, environmental, and genetic variation. Through differentiation, cells acquire highly specialized functions, but need to still maintain their general abilities to accurately regulate both essential pathways as well as responses to changes in the environment. To achieve robustness, regulatory processes must be capable of attenuating expression variability of essential genes (Bartha et al., 2018), while still allowing, or possibly amplifying (Eldar and Elowitz, 2010; Urban and Johnston, 2018), variability in expression for genes that are required for differentiation or responses to environmental changes and external cues. How cells can achieve such precision and robustness remains elusive.

Genetic variation affects the expression level (Montgomery et al., 2010; Pickrell et al., 2010; Stranger et al., 2007) of the majority of human genes (GTEx Consortium, 2017; Lappalainen et al., 2013; Storey et al., 2007). However, genes are associated with highly different effect sizes, with ubiquitously expressed or essential genes frequently being less affected (GTEx Consortium, 2017). This indicates that genes associated with different regulatory programs are connected with different mechanisms or effects of mutational robustness (Payne and Wagner, 2015). Multiple transcription factor (TF) binding sites may act to buffer the effects of mutations in promoters (Spivakov et al., 2012), and promoters can have highly flexible transcription start site (TSS) architectures (Akalin et al., 2009; Carninci et al., 2006; Lehner, 2008). This demonstrates that the sequence and architecture of a promoter may influence its variability in expression across individuals.

Previous studies aimed at identifying processes involved in the regulation of gene expression variability have indeed revealed regulatory features mostly associated with the promoters of genes, such as CpG islands and TATA-boxes (Morgan and Marioni, 2018; Ravarani et al., 2016; Sigalova et al., 2020), the chromatin state around gene TSSs (Faure et al., 2017), and the propensity of RNA polymerase II to pause downstream of the TSS (Boettiger and Levine, 2009). These studies have relied on model organisms or focused on transcriptional noise across single cells. As of yet, regulatory features have not been thoroughly studied from the perspective of variability in promoter activity or across human individuals. Furthermore, it is unclear if regulation of variability mainly acts to attenuate variability to achieve stable expression for certain genes or if independent regulatory processes act in parallel to amplify variability for other genes.

Here, we provide a comprehensive characterization of the sequences, TSS architectures, and regulatory processes determining variability of promoter activity across human lymphoblastoid cell lines (LCLs). We find that variability in promoter activity is to a large degree reflected by the promoter sequence, notwithstanding possible genotypic differences. Furthermore, the presence of binding sites for specific TFs, including those of the ETS family, are highly predictive of low promoter variability independently of their impact on promoter expression levels. In addition, we demonstrate that differences in the variability of promoters reflect their involvement in distinct biological processes, indicating a selective tradeoff between stability and plasticity of promoters. Finally, we show that flexibility in TSS usage is associated with attenuated promoter variability. Our results reveal a novel mechanism that confers mutational robustness to gene promoters via switches between proximal core promoters. This study provides fundamental insights into transcriptional regulation, indicating shared mechanisms that can buffer stochastic, environmental, and genetic variation and how these affect the responsiveness and cell-type restricted activity of genes.

## Results

### TSS profiling reveals variability in promoter activity across individuals

To characterize human variability in promoter activities, we profiled TSSs using CAGE (Cap Analysis of Gene Expression (Takahashi et al., 2012); Fig. 1A) across 108 Epstein-Barr virus (EBV)-transformed LCLs (Auton et al., 2015) of African origin, 89 from Yoruba in Ibadan, Nigeria (YRI) and 19 from Luhya in Webuye, Kenya (LWK) (Supplementary Table 1). The samples had a balanced sex ratio, 56 females and 52 males, and no observable population stratification in the expression data (Supplementary Fig. 1). With CAGE, TSSs can be mapped with single base pair resolution and the relative number of sequencing reads supporting each TSS gives simultaneously an accurate estimate of the abundance of its associated RNA (Kawaji et al., 2014). The CAGE data across the LCL panel therefore give us a unique opportunity to both estimate variability in promoter activity and characterize the regulatory features influencing such variability.

**Figure 1:**
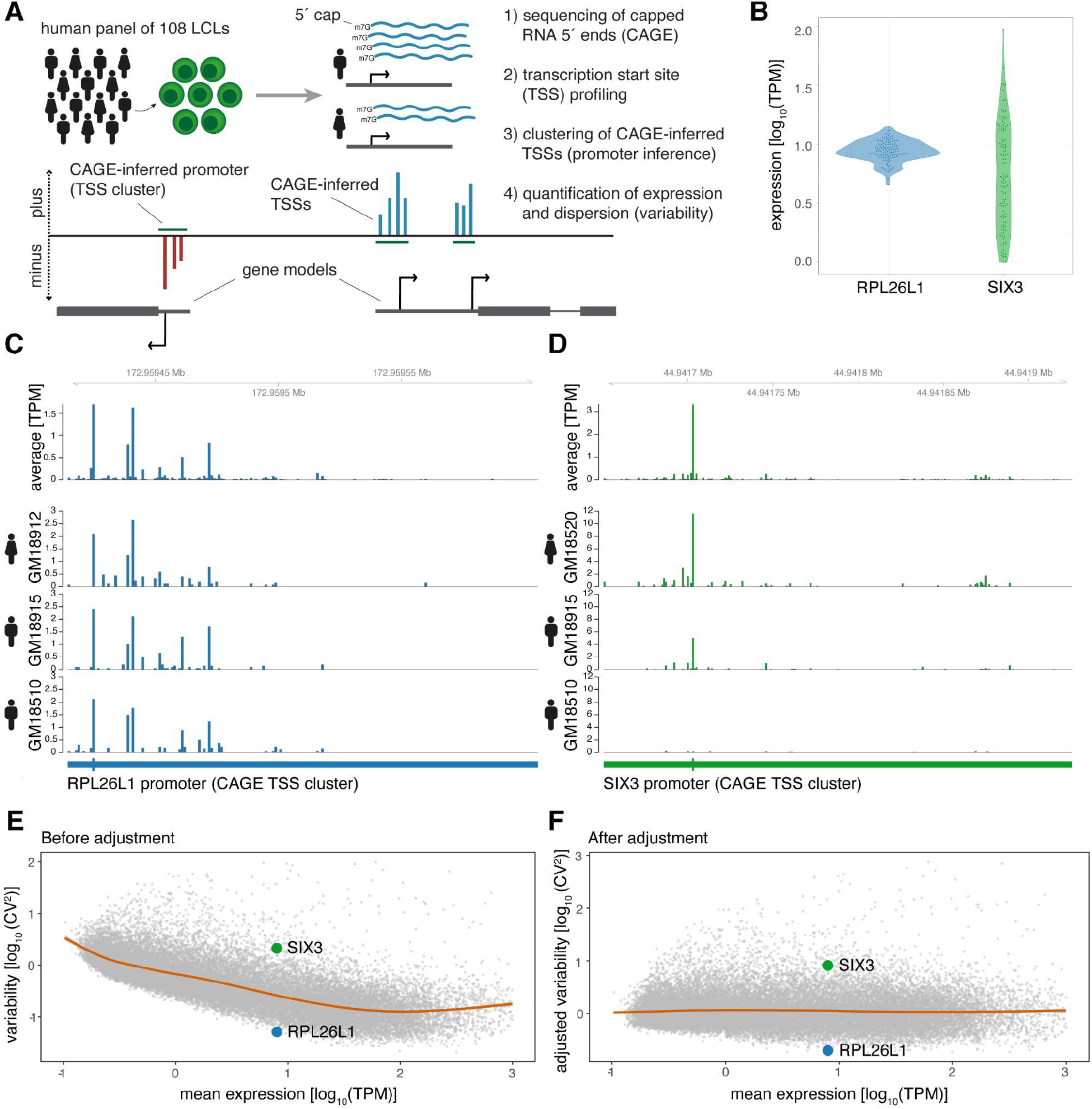
CAGE profiling of TSSs reveals diverse promoter variability across individuals. **A**: Illustration of the experimental design and approach for measuring promoter activity and variability. Capped 5’ ends of RNAs from LCLs derived from 108 individuals were sequenced with CAGE, followed by individual-agnostic positional clustering of proximal CAGE-inferred TSSs (first 5’ end bp of CAGE reads). The expression level of the resulting CAGE-inferred promoters proximal to annotated gene TSSs were quantified in each individual and used to measure promoter variability. **B**: Example of promoter activity (TPM normalized count of CAGE reads) across individuals for a low variable promoter (gene *RPL26L1*) and a highly variable promoter (gene *SIX3*) with similar average expression across the panel. **C-D**: Genome tracks for two promoters showing average TPM-normalized CAGE data (expression of CAGE-inferred TSSs) across individuals (top track) and TPM-normalized CAGE data for three individuals (bottom tracks) for a low variable promoter (panel C, gene *RPL26L1*) and a highly variable promoter (panel D, gene *SIX3*). **E-F**: The CV^2^ (squared coefficient of variation) and mean expression relationship of 29,001 CAGE-inferred promoters across 108 individuals before (E) and after (F) adjustment of the mean expression-dispersion relationship. The CV^2^ and mean expression are log_10_ transformed, orange lines show loess regression lines fitting the dispersion to the mean expression level, and example gene promoters from B-D are highlighted in colors.

We identified 29,001 active promoters of 15,994 annotated genes (Frankish et al., 2019) through positional clustering of proximal CAGE-inferred TSSs on the same strand (Fig. 1A) (Carninci et al., 2006) detected in at least 10% of individuals (Supplementary Table 2). This individual-agnostic strategy ensured a focus on promoters that are active across multiple individuals while also allowing for the measurement of variability in promoter activity across the panel. For example, the CAGE data revealed that the promoters of gene *RPL26L1*, encoding a putative component of the large 60S subunit of the ribosome, and transcription factor gene *SIX3* have highly different variance yet similar mean expression across individuals (Fig. 1B-D).

We used the squared coefficient of variation (CV^2^) as a measure of promoter expression dispersion, revealing how the normalized expression across individuals deviates from the mean for each identified promoter. We observed that the promoter CV^2^ decreases by increasing mean expression (Fig. 1E) (Eling et al., 2018; Kolodziejczyk et al., 2015; Sigalova et al., 2020). To account for this bias, we subtracted the expected dispersion for each promoter according to its expression level (Kolodziejczyk et al., 2015; Newman et al., 2006). Importantly, rank differences in promoter dispersion were maintained for each expression level after adjustment, as seen for promoters of genes *RPL26L1* and *SIX3* (Fig. 1E,F). This strategy thus allowed us to investigate how promoter architecture and sequence determine variability in promoter activity across the panel separately from its impact on expression level (Fig. 1F).

### Promoter expression variability is reflected by the promoter sequence

To investigate if local sequence features at promoters reflect their variability in activity, we applied machine learning (convolutional neural network (CNN); Supplementary Fig. 2A; see Methods) to discern low variable promoters (N=5,054) from highly variable promoters (N=5,683) based on their local DNA sequence alone. We considered the genomic reference sequence to model the intrinsic component of variability encoded within the promoter sequence independently of local genetic variants within the panel. The resulting model was capable of distinguishing between these promoter classes with high accuracy (area under receiving operating curve (AUC)=0.84 for the out of sample test set; Supplementary Fig. 2B), equally well for highly and low variable promoters (per-class test set F1 scores of 0.76 and 0.77, respectively).

To assess which sequence features the CNN had learned to distinguish the classes, we calculated importance scores using DeepLift (Shrikumar et al., 2019) for each nucleotide in the input sequences for predicting low and high promoter variability. This approach relies on backpropagation of the contributions of all neurons in the CNN to the input features, nucleotides, and can therefore be used to identify properties or short stretches of DNA indicative of amplifying or attenuating expression variability. We applied motif discovery on clustered stretches, so called metaclusters, of the input sequences with high importance scores (Shrikumar et al., 2020) and matched the identified metaclusters to known TF binding motifs (Fornes et al., 2020). This strategy revealed TFs indicative of either high or low promoter variability (Fig. 2A-C). Noteworthy, we observed motifs for the ETS superfamily of TFs, including ELK1, ETV6, and ELK3, associated with low variable promoters, and motifs for PTF1A, ASCL2, and FOS-JUN heterodimer (AP-1) among highly variable promoters. These results demonstrate that the promoter sequence and its putative TF binding sites are predictive of the expression variability of a promoter.

**Figure 2:**
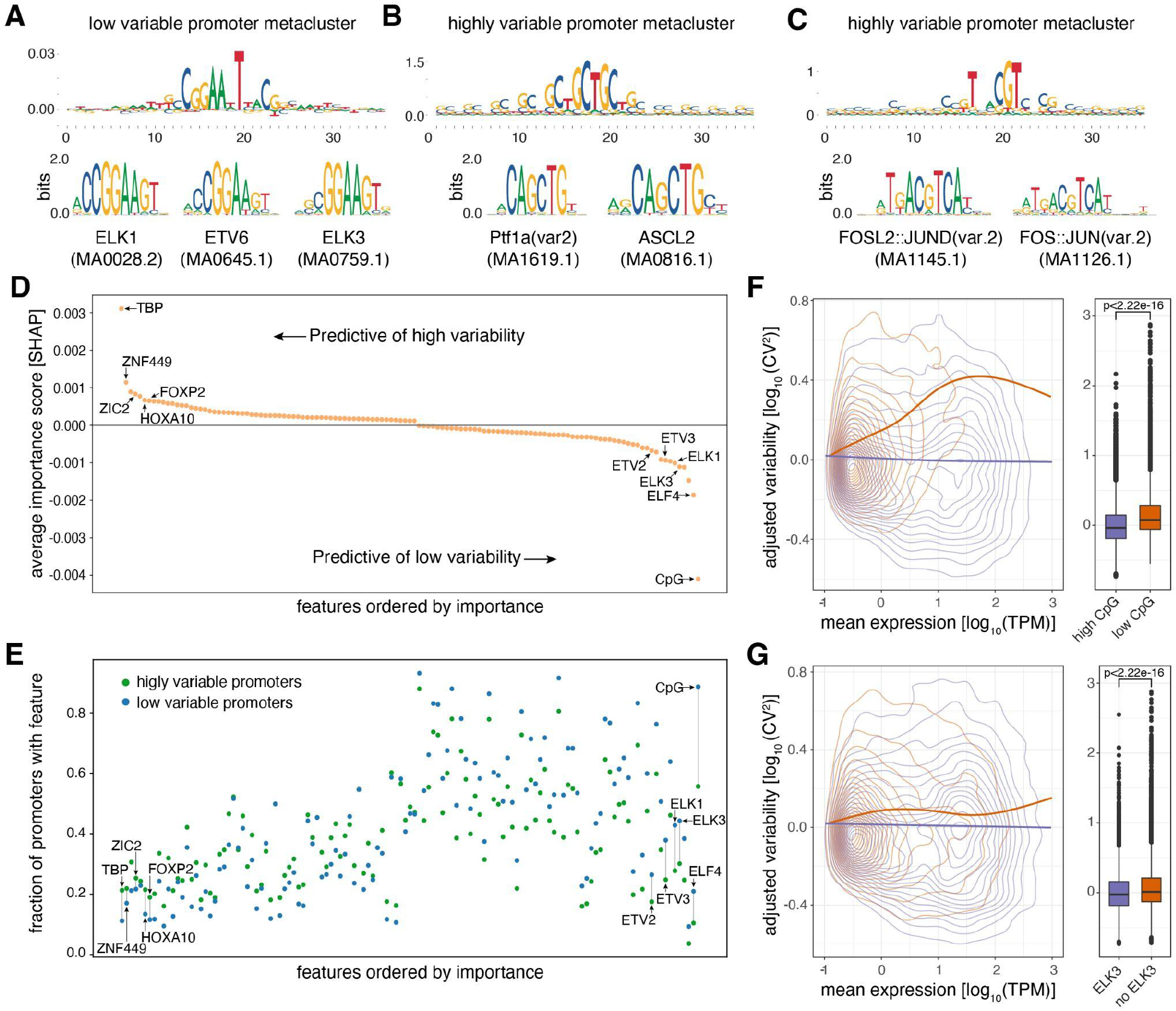
Promoter sequence features are highly predictive of promoter variability. **A**: Sequence logo of a metacluster (top) identified for low variable promoter sequences that matches known TF motifs (bottom) for ETS factors ELK1, ETV6, and ELK3. **B-C**: Sequence logos of two metaclusters (top) identified for highly variable promoter sequences that match known TF motifs (bottom) for PTF1A and ASCL2 (B) and FOSL2-JUND and FOS-JUN heterodimers (C). **D**: Average contribution (SHAP values) of CpG content and each of the 124 TFs identified as important for predicting promoter variability. Features are ordered by their average contribution to the prediction of highly variable promoters and selected TFs are highlighted. For a full version of the plot see Supplementary Figure 4A. **E**: The frequency of predicted TF binding sites (presence/absence) in highly variable (green) and low variable (blue) promoters. TFs are ordered as in D. For a full version of the plot see Supplementary Figure 4B,C. **F-G**: Promoters split into groups based on the presence/absence of high CpG content (F), and predicted binding sites of ELK3 (G). For both features displayed in panels F and G, the left subpanels display the relationship between log_10_-transformed mean expression levels and adjusted log_10_-transformed CV^2^ with loess regression lines shown separately for each promoter group. The right subpanels display box-and-whisker plots of the differences in adjusted log_10_-transformed CV^2^ between the two promoter groups (central band: median; boundaries: first and third quartiles; whiskers: +/− 1.5 IQR). P-values were determined using the Wilcoxon rank-sum test (***: p<0.05).

### Sequence features of promoters are highly predictive of promoter variability

To systematically test how predictive TF binding sites are of the variability of active promoters, we made use of binding sites predicted from motif scanning for 746 TFs (Fornes et al., 2020). TF binding site profiles and low/high CpG content (Supplementary Fig. 3A) were collected for each identified promoter and the resulting feature data were used to train a machine learning (random forest) classifier features associated with either high or low variability (low variable N=5,054, highly variable N=5,683). Feature selection (Kursa and Rudnicki, 2010) identified 124 of the 746 TFs as well as CpG ratio to be important for classification, and a classifier based on these selected features demonstrated high predictive performance (AUC=0.79; per-class F1 score of 0.73 and 0.68 for low and highly variable promoters, respectively; Supplementary Fig. 3B), reinforcing the strong link observed between DNA sequence and promoter variability (Supplementary Fig. 2B).

Reverse engineering of the random forest classifier (SHAP, Shapley additive explanations) (Lundberg and Lee, 2017) allowed us to calculate the marginal contribution of each of the 125 selected features to the prediction of variability class for each promoter and whether the feature on average is indicative of amplifying or attenuating variability of expression when present in the promoter sequence (Fig. 2D; Supplementary Fig. 4A). We identified the presence of high observed/expected CpG ratio and TATA-binding protein (TBP) binding sites (TATA-boxes) to be the strongest predictive features of low and high promoter variability, respectively. Although the remaining TFs contribute only marginally on their own to the predictions compared to TATA-box and high CpG content status (Fig. 2D; Supplementary Fig. 4A), a baseline model (decision tree) based on CpG ratio and TBP binding site presence alone yielded worse performance than the full model (AUC=0.71 versus 0.79 for the baseline model and the full model, respectively; Supplementary Fig. 3B). This demonstrates that the TF binding grammar contributes to a promoter’s expression variability.

TFs associated with highly variable promoters are mostly related to tissue specific or developmental regulation (e.g., FOXP2, HOXA10) while TFs predictive of low promoter variability are generally associated with ubiquitous activity across cell types and a diverse range of basic cellular processes (e.g., ELK1, ELF4, ETV3). In addition, TFs predictive of high variability (e.g., ZIC2, ZNF449, HOXA10) tend to have binding sites in relatively few highly variable promoters while TFs predictive of low promoter variability (e.g., ELK1, ELK3) show a propensity for having binding sites present in a large number of promoters (Fig. 2E; Supplementary Fig. 4B,C). This suggests that variably expressed promoters have diverse TF binding profiles and that the regulatory grammar for promoter stability is less complex.

Although the adjusted dispersion of promoters was separated from their expression level (Fig. 1E), we observed that the presence of binding sites for some TFs that are predictive of promoter variability are also associated with promoter expression level (Supplementary Fig. 5). Importantly, despite this association, the effect of identified features on promoter variability is still evident across a range of promoter expression levels (Fig. 2F,G). This is particularly apparent for CpG islands, which seem to have an attenuating effect on promoter variability regardless of expression level (Fig. 2F).

Many of the TFs identified as being predictive of low variability (e.g., ELK1, ELK3, ELF4, ETV2, ETV3) belong to the ETS family of TFs (Fig. 2D; Supplementary Fig. 4A), a large group of TFs that are conserved across Metazoa and characterized by their shared ETS domain that binds 5′-GGA(A/T)-3′ DNA sequences (Sharrocks, 2001). ETS factors are important regulators of promoter activities in lymphoid cells (Hepkema et al., 2020), but are generally involved in a wide range of crucial cellular processes such as growth, proliferation, apoptosis, and cellular homeostasis (Kar and Gutierrez-Hartmann, 2013; Oikawa and Yamada, 2003; Suico et al., 2017). Furthermore, different ETS factors can bind in a redundant manner to the same promoters of housekeeping genes (Hollenhorst et al., 2011, 2007). However, the shared DNA binding domain of ETS factors makes it hard to discern individual factors based on their binding motifs alone (Fig. 2A). Although in general linked with higher promoter activity (Curina et al., 2017), ETS binding site presence is associated with lower variability across all expression levels (Fig. 2G). In addition, the degree of promoter variability decreases by an increasing number of non-overlapping ETS binding sites (Supplementary Fig. 6A), regardless of promoter expression level (Supplementary Fig. 6B), suggesting that multiple ETS binding sites can either facilitate cooperativity between ETS factors or provide robustness to stabilize promoter variability across individuals.

Taken together, our results indicate that promoter sequence can influence both low and high promoter variability across human individuals independently from its impact on expression level. Our results further indicate that variable promoters exhibit highly diverse binding grammars for TFs that are associated with relatively few promoters, while a more uniform regulatory grammar is indicated for stable promoters, being highly associated with higher CpG content and ETS binding sites.

### Variability in promoter activity reflects plasticity and robustness for distinct biological functions

The high performance of predicting promoter variability from local DNA sequence and the distinct TF binding profiles of low and highly variable promoters implies that attenuation and amplification of variability are driven by distinct regulatory mechanisms. This argues that favoring robustness (low variability) over plasticity (high variability) should reflect the biological processes where this provides regulatory advantages. Supporting this hypothesis, we observed that low variable promoters were highly enriched with basic cellular housekeeping processes, in particular metabolic processes (Fig. 3A). In contrast, highly variable promoters were enriched with more dynamic biological functions, including signaling, response to stimulus, and developmental processes.

**Figure 3:**
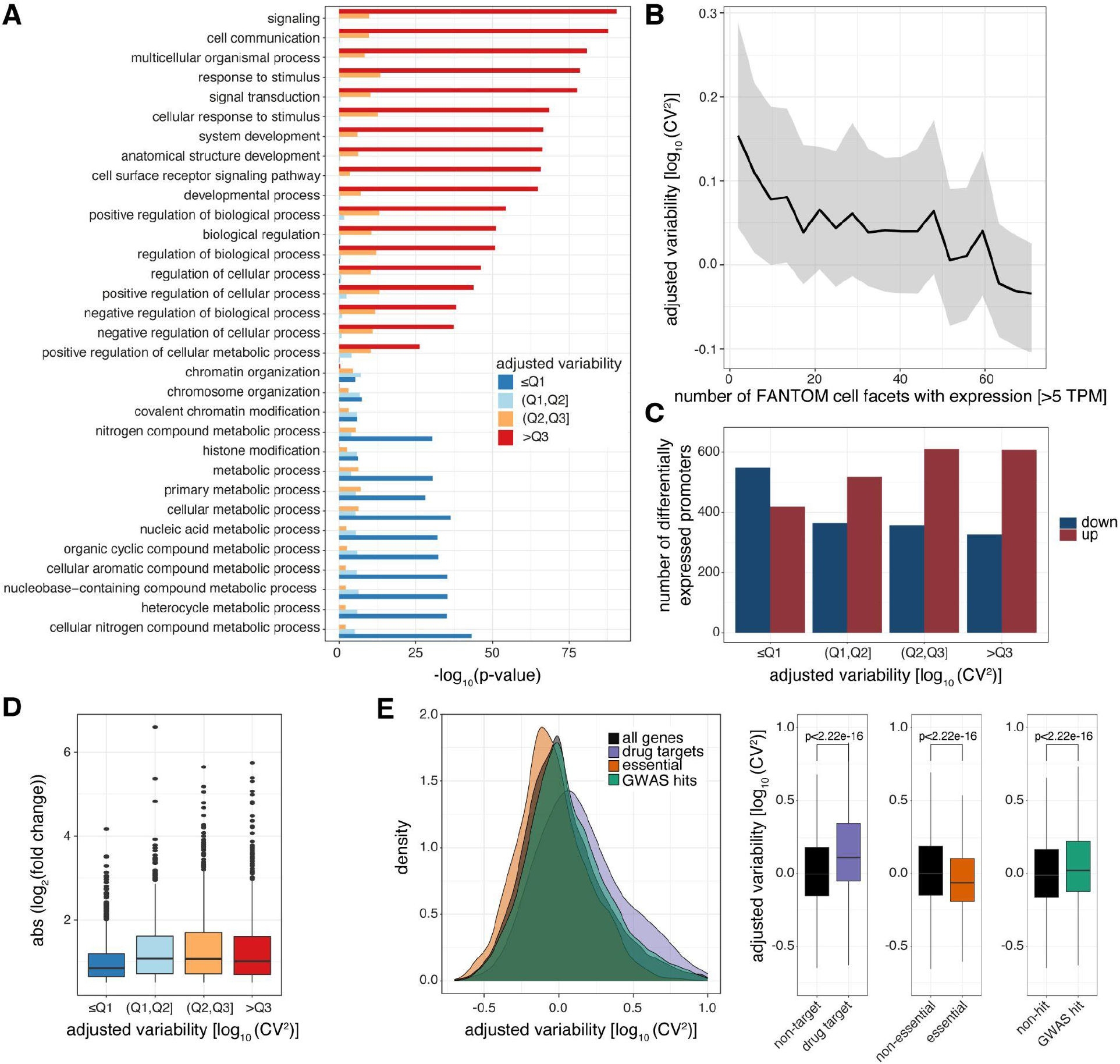
Levels of promoter variability are reflective of distinct biological processes and a selective trade-off between robustness and plasticity. **A**: GO term enrichment, for biological processes, of genes split by associated promoter variability quartiles (Q1, Q2, Q3). Top 10 GO terms of all groups are displayed and ranked based on p-values of the >Q3 variability group. **B**: Median promoter variability (line) and interquartile range (shading), as a function of the number of FANTOM cell facets (grouping of FANTOM CAGE libraries associated with the same Cell Ontology term) that the associated gene is expressed in (mean facet expression >5 TPM). **C**: The number of differentially expressed promoters, split by variability quartiles, after 6h TNFα treatment. Promoters are separated into down-regulated (blue) and up-regulated (red). P-values were calculated using Fisher’s exact test. **D**: Absolute log2 fold change of differentially expressed promoters, split by variability quartiles, after 6h of TNFα treatment. **E**: Distribution of promoter variability associated with drug-targets (purple), essential (orange), or GWAS hits (green) genes, compared to all promoters (black). Left: density plot of promoter variability per gene category. Right: Box-and-whisker plots of promoter variability split by each category of genes. P-values were determined using the Wilcoxon rank-sum test. For all box-and-whisker plots, central band: median; boundaries: first and third quartiles; whiskers: +/− 1.5 IQR.

Interestingly, the same features found to be predictive of low and high promoter variability across individuals, including CpG-content and TATA-boxes (TBP binding sites), are also associated with low and high transcriptional noise across individual cells (Faure et al., 2017; Morgan and Marioni, 2018). The presence of a TATA-box is also associated with high gene expression variability in flies (Sigalova et al., 2020). This suggests that some of the same underlying regulatory mechanisms that dictate low or high transcriptional noise across single cells are maintained and conserved between humans and flies at an individual level and manifested to control low and high expression variability across a population, respectively, as well as housekeeping or restricted activity across cell types.

In agreement, genes of highly variable promoters tend to have higher transcriptional noise than those of low variable ones across GM12878 single cells (Cohen’s d=0.411, p<2.2×10^−16^, two sample t-test; Supplementary Fig. 7A; Supplementary Table 3). Furthermore, we observed an inverse correlation between variability in promoter activity and the number of cell types (FANTOM Consortium and the RIKEN PMI and CLST (DGT) et al., 2014) and tissues (GTEx Consortium, 2017) the corresponding gene is expressed in (Spearman’s rank correlation ρ = −0.21 and −0.15 for cell types and tissues, respectively, p< 2.2×10^−16^; Fig. 3B; Supplementary Fig. 7B), demonstrating that highly variable promoters are more cell-type and tissue specific in their expression.

The restricted expression (Supplementary Fig. 7B), biological processes (Fig. 3A), and promoter TF grammar (Fig. 2D,E) of genes associated with highly variable promoters led us to hypothesize that these are more prone to respond to external stimuli. Tumor necrosis factor (TNFα) induces an acute and time-limited gene response to NFkB signaling (Nelson et al., 2004; Turner et al., 2010), with negligible impact on chromatin topology (Jin et al., 2013), and is therefore suitable to study gene responsiveness. We profiled GM12878 TSSs and promoter activities with CAGE before and after 6 hours treatment with TNFα (Supplementary Table 4). This revealed an enrichment of up-regulated promoters among highly variable promoters (odds ratio (OR)=1.529, p=4.563×10^−8^, Fisher’s exact test) as posited, while low variable promoters were mostly unaffected or down-regulated (OR=0.459, p<2.2×10^−16^, Fisher’s exact test; Fig. 3C). In addition, low variable promoters had a weaker response (Fig. 3D).

Furthermore, we observed drug-target genes and genes with GWAS hits to be regulated by highly variable promoters but essential genes to be regulated by low variable promoters (Fig. 3E). In contrast, when we compared promoter expression between these same groups of genes we observed no association with drug-targets or GWAS-associated genes. Although essential genes are associated with higher promoter expression, this association is comparably weaker than that with promoter variability (Supplementary Fig. 7C).

Taken together, our results demonstrate the importance of low promoter variability for cell viability and survival in different conditions and reveal the responsiveness of highly variable promoters. They further indicate that the variability observed in promoter activity across individuals is strongly associated with the regulation of its associated gene, the expression breadth across cell types, and to some extent also the transcriptional noise across single cells, which reflects a selective trade-off between high stability and high responsiveness and specificity.

### Promoters with low variability have flexible transcription initiation architectures

Promoters are associated with different levels of spread of their TSS locations, which has led to their classification into broad or narrow (sharp) promoters according to their positional width (Akalin et al., 2009; Carninci et al., 2006; Lehner, 2008). Although the shape and distinct biological mechanisms of these promoter classes, e.g., housekeeping activities of broad promoters, are conserved across species (Carninci et al., 2006; Hoskins et al., 2011), the necessity for positional dispersion of TSSs and its association with promoter variability are poorly understood.

Analysis of promoter widths revealed only a weak relationship with promoter variability. We observed an enrichment of highly variable promoters within narrow promoters having an interquartile range (IQR) of their CAGE-inferred TSSs within a width of 1 to 5 bp (p<2.2×10^−16^, OR=2.04, Fisher’s exact test). Low variable promoters, on the other hand, were enriched among those of size 10 to 25 bp (p<2.2×10^−16^, OR=1.44, Fisher’s exact test), but beyond this width the association is lost (Supplementary Fig. 8A). To simultaneously capture the spread of TSSs and their relative frequencies compared to total RNA expression within a promoter, we considered a width-normalized Shannon entropy as a measure of TSS positional dispersion (Hoskins et al., 2011). This measure will discern promoters whose relative TSS expression is concentrated to a small subset of their widths (low entropy) from those with a more even spread (high entropy). We observed that low variable promoters are associated with a higher entropy than promoters with high variability (Fig. 4A). Consistently, low variable promoters tend to have more TSSs substantially contributing to their overall expression across individuals (Supplementary Fig. 8B). We reasoned that a weaker association between low promoter variability and broad width than with high entropy may be due to low variable promoters being composed of multiple clusters of TSSs (multi-modal peaks) from independent core promoters. Indeed, decomposition of multi-modal peaks within the CAGE TSS signals of promoters (Supplementary Table 5) demonstrated that higher entropy reflects an increased number of decomposed promoters, as indicated by their number of local maxima of CAGE signals (Fig. 4B).

**Figure 4:**
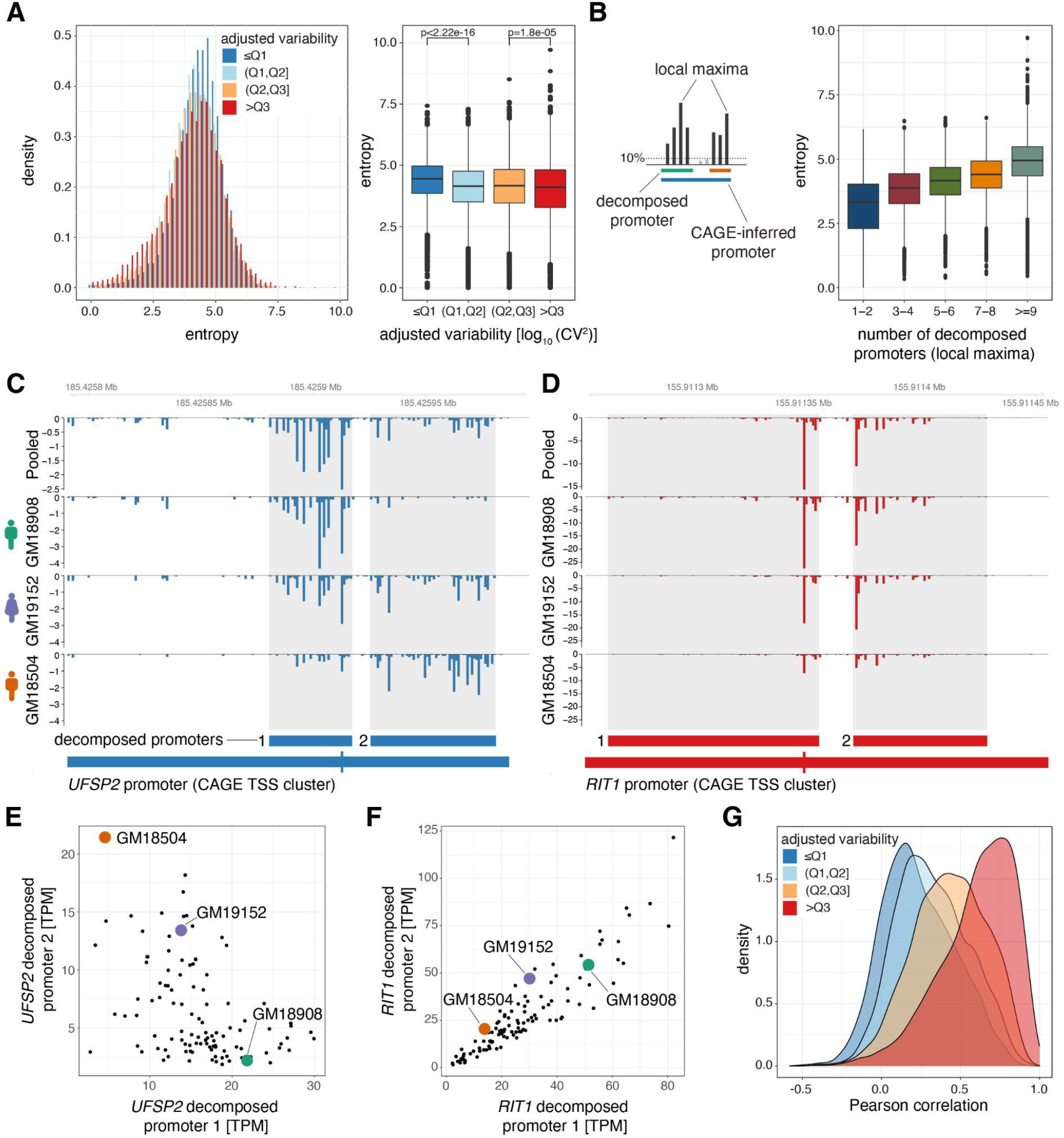
Low variable promoters exhibit flexibility in transcription initiation architecture. **A**: Promoter shape entropy for promoters split by variability quartiles, displayed as densities (left) and in a box-and-whisker plot (right). **B**: Illustration of the local maxima decomposition approach (left; see Materials and Methods) and box-and-whisker plot displaying the relationship between the Shannon entropy and the number of local maxima-inferred decomposed promoters. **C-D**: Examples of two promoters each containing two decomposed promoters exhibiting low correlation across individuals (panel C, gene UFSP2) and high correlation across individuals (panel D, gene RIT1). Both panels display genome tracks of average, TPM-normalized CAGE-inferred TSS expression levels across the panel (Pooled, top track) and for three individuals (GM18908, GM19152, GM18504, lower tracks). Below the genome tracks, the original promoter and resulting decomposed promoters (shaded in genome tracks) are shown. **E-F**: Relationship between TPM-normalized CAGE expression of decomposed promoter 1 (x-axis) and 2 (y-axis) across all 108 LCLs for example genes *UFSP2* (E) and *RIT1* (F). The expression values for individuals included in panels B and C are highlighted. **G**: Densities of the lowest Pearson correlation between all pairs of decomposed promoters originating from the same promoter across all CAGE-inferred promoters with at least two decomposed promoters. For all box-and-whisker plots, central band: median; boundaries: first and third quartiles; whiskers: +/− 1.5 IQR.

The decomposed promoters of gene *UFSP2* (Fig. 4C,E) clearly illustrate that the activity of sub-clusters of TSSs within promoters and their contributions to the overall activity of the encompassing promoter can vary to a great extent between individuals. In contrast, the decomposed promoters of gene *RIT1* (Fig. 4D,F) contribute equally to the overall activity of the encompassing promoter across individuals. To assess in general how individual decomposed promoters influence the overall promoter variability, we calculated the expression-adjusted dispersion (adjusted log_10_-transformed CV^2^) of local-maxima decomposed promoters. Interestingly, many of the decomposed promoters showed a vastly different variability across individuals compared to the promoters they originate from (Supplementary Fig. 8C). This disagreement indicates that decomposed promoters within the same promoter reflect core promoters that may either operate and be regulated independently of each other or differ in their ability to compete for the transcriptional machinery, both of which may contribute to the overall robustness or plasticity of the promoter and, in turn, the gene. As highly multimodal peaks are mainly found to be associated with low variable promoters, we hypothesized that this flexibility in core promoter usage may act as a compensatory mechanism to stabilize their expression.

If the effects of significant changes in expression across the panel are masked by compensatory changes in decomposed promoter usage within the same promoter, this would be revealed by low or even negative expression correlation between decomposed promoters (e.g., decomposed promoters 1 and 2 of *UFSP2*, Fig. 4C,E). Indeed, we observed a strong association between promoter variability and the minimal expression correlation between decomposed promoter pairs within a promoter (Fig. 4G). Low variable multi-modal promoters are associated with weakly or even negatively correlated pairs of decomposed promoters. In contrast, highly variable multi-modal promoters are associated with moderately or highly correlated pairs of decomposed promoters. The weak expression correlation between decomposed promoters of low variable promoters demonstrates that decomposed promoters may operate independently of each other, while negatively correlated pairs indicate a competition for the transcriptional machinery or a compensatory shift between decomposed promoters. The association between decomposed promoter correlation and overall promoter stability was maintained when all decomposed promoter pairs were considered (Supplementary Fig. 8D), and could not be explained by CpG island status (Supplementary Fig. 8E). However, decomposed promoter expression correlation was associated with promoter width (Supplementary Fig. 8F), demonstrating that complex low variable promoters with multiple decomposed promoters require larger promoter width, while broad promoter width does not necessarily lead to lower promoter variability.

The spread and dominant position of TSSs in broad promoters are tightly linked to immediate downstream (+1) nucleosome positioning, and changes in +1 nucleosome positioning can alter the preferred TSS (Dreos et al., 2016; FANTOM Consortium and the RIKEN PMI and CLST (DGT) et al., 2014; Haberle et al., 2014). Hence, the variability and multi-modal TSS patterns of promoters could be related to their nucleosomal architectures. Indeed, comparison of +1 nucleosome positioning relative to the dominant TSS position of promoters across the panel revealed that low variable promoters tend to have stronger +1 nucleosome positioning (Supplementary Fig. 9A,B). However, when analyzing specifically multi-modal low variable promoters (containing at least two decomposed promoters), a strong +1 nucleosome was only observed for promoters with highly correlated decomposed promoters (Supplementary Fig. 9C,D). Furthermore, highly variable multi-modal promoters and those containing low correlated pairs of decomposed promoters exhibited more fuzzy +1 nucleosome positioning across cells (Supplementary Fig. 9E,F). Our results thus demonstrate that low variable promoters with flexible TSS usage, i.e., having weakly or negatively correlated decomposed promoters, are characterized by less restrictive and more fuzzy +1 nucleosome positioning.

Taken together, our results demonstrate that flexible usage of core promoters within promoters with permissive nucleosomal architectures provide stability to the overall expression of a large subset of gene promoters with low variability.

### Alternative TSSs of low variability promoters indicate a novel mechanism of mutational robustness

While genetic variants associated with gene expression levels (expression quantitative trait loci, eQTLs) frequently occur within gene promoters, they are rarely found associated with housekeeping or ubiquitously expressed genes, and when they are, they have limited effect sizes (GTEx Consortium, 2017). One explanation for this observation is that mutations that would significantly alter the expression of such genes would be detrimental to cell viability. In addition, the rare and limited effects of eQTLs on housekeeping genes might be due to mechanisms promoting mutational robustness. Our results (Fig. 4) indicate that a flexible TSS architecture within a promoter may provide such a mechanism and thereby mask the effects genetic variants may have on individual decomposed promoters.

To test if flexibility in core promoter usage within a promoter may cause mutational robustness, we first performed local eQTL analysis on promoters (within 25kb). We tested both the association between the genotypes of common genetic variants (MAF ≥ 10%) and the expression of promoters (promoter eQTL, prQTLs; Fig. 5A, top) as well as those of decomposed promoters (decomposed promoter eQTL, dprQTL). 2,457 promoters were associated with at least one prQTLs (5% FDR; Supplementary Table 6). While prQTLs were observed across all levels of promoter variability, they were more commonly associated with highly variable promoters (Fig. 5B). Fewer prQTL single nucleotide polymorphisms (SNPs) and, in general, common variants were found proximal to low variable promoters, indicating a negative selection for these. As expected, the effect size for the most significant prQTL variant (lead SNP) for each promoter was positively associated with the expression variability of the promoter (Spearman’s rank correlation ρ = 0.16, p< 2.2×10^−16^, Supplementary Fig. 10A). This indicates that, in addition to having fewer proximal genetic variants, low variable promoters are less likely to have prQTLs with large regulatory effects. However, QTLs of decomposed promoter expression (dprQTLs) exhibited similar prevalence (Supplementary Fig. 10C) and maximum effect sizes (Supplementary Fig. 10D) across promoter variability classes. This suggests that a flexible TSS architecture can limit the impact of genetic variants on promoter expression.

**Figure 5:**
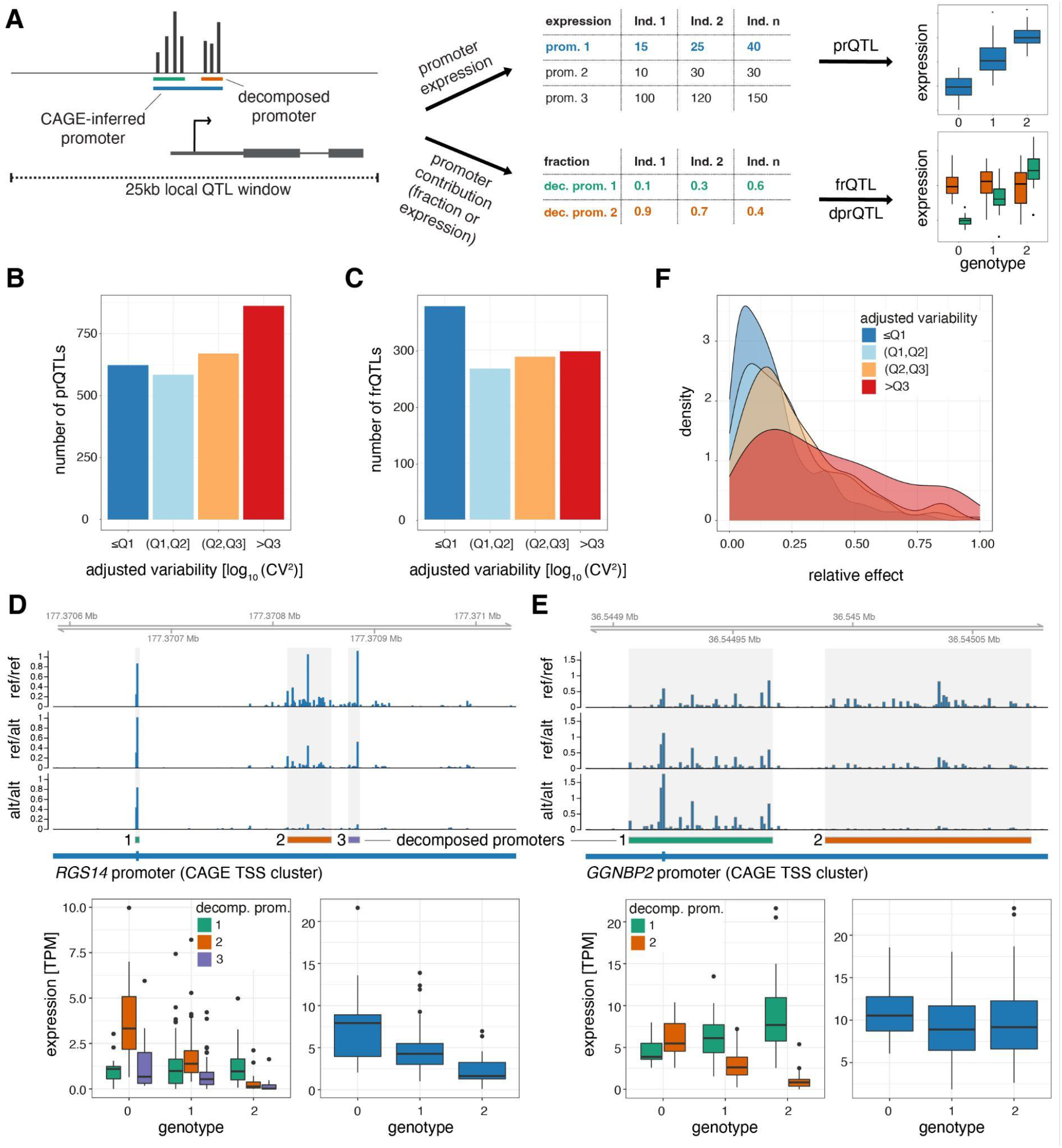
Plasticity in TSS usage is linked with increased mutational robustness. **A**: Illustration of the strategy for testing the effects of genetic variants on promoter expression (prQTLs, TPM-normalized CAGE counts), decomposed promoter expression (dprQTLs, TPM-normalized CAGE counts), and decomposed promoter contribution to the encompassing promoter expression (frQTLs, ratios of TPM-normalized CAGE counts between decomposed and encompassing promoters). For both approaches only SNPs within 25kb of the promoter CAGE signal summit were tested. **B**: Number of prQTLs detected (FDR<0.05), split by promoter variability quartiles. **C**: Number of encompassing promoters with at least one frQTL detected for a contained decomposed promoter (FDR<0.05), split by encompassing promoter variability quartiles. **D-E**: Examples of two promoters associated with frQTLs for a highly variable promoter with limited buffering of promoter expression (panel D, gene *RGS14*) and for a low variable promoter with strong buffering of promoter expression (panel E, gene *GGNBP2*). Upper panels display genome tracks showing average TPM-normalized CAGE data across homozygous individuals for the reference allele (top track), heterozygous individuals (middle track), and homozygous individuals for the variant (alternative) allele (bottom track). The bottom left subpanels display box-and-whisker plots of the differences in TPM-normalized CAGE data between genotypes for each decomposed promoter. The bottom right subpanels display box-and-whisker plots of the differences in TPM-normalized CAGE data between the three genotypes for the original encompassing promoter. For all box-and-whisker plots, central band: median; boundaries: first and third quartiles; whiskers: +/− 1.5 IQR. **F**: Density plot of the maximal relative change in expression between reference and variant alleles (relative effect size) for the most significant frQTL of each broad promoter with FDR ≥ 5%, split by variability quartiles.

To further investigate the observed disagreement between the variability of promoters and their decomposed promoters, we tested the association between the genotypes of common genetic variants and the contribution of decomposed promoters to the overall expression of their encompassing (non-decomposed) promoters (fraction eQTL, frQTL; Fig. 5A, bottom). We identified 1,230 promoters to be associated with at least one frQTL (5% FDR; Supplementary Table 7). Unlike the prQTLs and dprQTLs, the frQTLs were more commonly associated with decomposed promoters from low variable promoters (Fig. 5C). Conceptually, the frQTLs can affect decomposed promoter usage and overall promoter expression levels to different degrees, as exemplified by the promoters of genes *RGS14* and *GGNBP2* (Fig. 5D,E). Gene *RGS14* has three decomposed promoters localized within its promoter (Fig. 5D), for which SNP rs56235845 (chr5:177371039 T/G) was strongly associated with the contribution to the overall promoter activity for only decomposed promoters 1 and 2 (frQTL beta=0.210, −0.181, −0.062; FDR=2.42×10^−5^, 2.54×10^−8^, 2.64×10^−2^, for decomposed promoters 1, 2, and 3, respectively). Despite the limited association of the variant with decomposed promoter 3, it still had a noticeable association with the overall promoter activity (prQTL beta=−2.47, FDR=3.57×10^−5^; Fig. 5D, bottom right). In contrast, SNP rs9906189 (chr17:36549567 G/A) was strongly associated with the contribution to the overall promoter activity for both decomposed promoters of gene *GGNBP2* (frQTL beta=0.222, −0.222; FDR=2.05×10^−26^, 2.05×10^−25^, for decomposed promoters 1 and 2, respectively), but in opposite directions (Fig. 5E). Interestingly, this switch in decomposed promoter usage translates into a limited effect on the overall promoter activity (prQTL beta=−0.063, FDR=0.989; Fig. 5E, bottom right).

Both examples, a partial shift (Fig. 5D) and a switch (Fig. 5E) in decomposed promoter usage, are indicative of plasticity in TSS usage, which can secure tolerable levels of steady-state mRNA. Although frQTLs were associated with promoters across the wide spectrum of promoter variabilities (Fig. 5C), they showed a large difference in their relative effect on the overall promoter activity (maximal relative change in expression between reference and variant alleles; Fig. 5F). frQTLs associated with highly variable promoters tend to have a larger relative effect on the overall promoter activity compared to frQTLs associated with low variable promoters. This association is further maintained at the gene level (adjusted RNA-seq (Lappalainen et al., 2013) CV^2^; Supplementary Fig. 10B; Supplementary Table 8), demonstrating that individual differences in decomposed promoter usage contribute to low promoter variability and, in turn, low gene variability. In total, we found 286 promoters (out of 1,230) of 284 genes to be associated with stabilizing frQTLs, for which the same SNP was associated with at least two decomposed promoters (5% FDR) with relative effects in opposite directions (Supplementary Table 9). Our results thus indicate that TSS usage flexibility confers mutational robustness that stabilizes the variability of promoters and their associated genes.

Taken together, integrating prQTLs, dprQTLs, and frQTLs provides novel insights into how common genetic variants can influence TSS usage in humans and its potential impact on gene expression. We demonstrate that low variable promoters characterized by multiple decomposed promoters (multi-modal TSS usage) are less affected by the presence of genetic variants compared to highly variable promoters. In addition, we find that part of this tolerance can be explained by a, previously unreported, mechanism of mutational robustness through plasticity in TSS usage. The prevalence of expression-independent decomposed promoters within low variable promoters, as suggested by low pairwise correlation, indicates an extensive regulatory role of TSS plasticity in attenuating expression variability of essential genes.

## Discussion

In this study we provide an extensive characterization of promoter-associated features influencing variability in promoter activity across human individuals and demonstrate their importance for determining stability, responsiveness, and specificity. Overall, we show that the local DNA sequence, putative TF binding sites, and transcription initiation architecture of promoters are highly predictive of promoter variability.

Although the classifier based on TF binding site sequence and CpG island status was able to predict promoter variability well (AUC=0.79 on the test set), it did not perform as well as the CNN model (AUC=0.84 on the test set), which was trained on DNA sequence alone. This indicates that additional information influencing variability may be encoded within the promoter sequence. For instance, the density and variations of Initiator elements within promoters could influence TSS flexibility (Carninci et al., 2006; Frith et al., 2007; Haberle et al., 2014; Nepal et al., 2020). In addition, di- or tri-nucleotide sequence patterns and stretches of high AT-richness, which influence local nucleosome positioning (Dreos et al., 2016; Haberle et al., 2014; Segal et al., 2006), impose different requirements for chromatin remodeling activities (Lorch et al., 2014) at gene promoters of low and high variability, which in turn may affect their variability and responsiveness. The promoter sequence may also encode a promoter’s intrinsic enhancer responsiveness (Arnold et al., 2017), which may influence its expression variability. Although current data cannot distinguish between direct or secondary effects, an increased variability mediated via enhancers is supported by a higher dependency on enhancer-promoter interactions for cell-type specific genes compared to housekeeping genes (Furlong and Levine, 2018; Schoenfelder and Fraser, 2019). However, compatibility differences between human promoter classes and enhancers only result in subtle effects *in vitro* (Bergman et al., 2022), suggesting that measurable promoter variability is likely a result of both intrinsic promoter variability and additive or synergistic contributions from enhancers. Directly modeling the influence and context-dependency of enhancers on promoter variability would therefore be important to further characterize regulatory features that may amplify gene expression variability.

Despite a clear association with high promoter CpG content and housekeeping genes, low variable promoters were not strongly associated with a broader width, which we would expect from promoters in CpG islands and with housekeeping activity (Carninci et al., 2006). Rather, our results suggest that low variability requires a certain minimum promoter width that can encompass a transcription initiation architecture competent of attenuating variability through flexible TSS usage. Switching between proximal clusters of TSSs (decomposed promoters) within a larger promoter is fundamentally different from that between alternative promoters (Garieri et al., 2017; Valen et al., 2008; Zhang et al., 2017), which will more likely lead to differences in transcript and protein isoforms. Rather, a flexible initiation architecture enables several points of entries for RNA polymerase II to initiate in the same promoter. This ensures proper gene expression across different cell types (FANTOM Consortium and the RIKEN PMI and CLST (DGT) et al., 2014; Kawaji et al., 2006) and developmental stages (Haberle et al., 2014). Interestingly, ETS factors, here associated with low variable promoters, can interact with transcriptional co-activators and chromatin modifying complexes (Curina et al., 2017; Göös et al., 2022). ETS factors may therefore play a role in TSS selection in promoters with multi-modal architectures (Lam et al., 2019). Here, we show that such flexibility is also associated with low variability across individuals for the same cell type. Our findings further indicate that plasticity in TSS usage within a promoter confers a, previously unreported, layer of mutational robustness that can buffer the effects of genetic variants, leading to limited or no impact on the overall promoter expression. Of note, the presence of weak or negatively correlated expression patterns between decomposed promoters for a large number of promoters suggests that such buffering events will be revealed for more genes with an increased sample size.

A flexibility in TSS usage may ensure transcriptional robustness of genes both in different environments and in the face of genetic variation. Since promoter shape is generally conserved across orthologous promoters (Carninci et al., 2006; Hoskins et al., 2011), it is plausible that robustness through flexible TSS usage is a conserved mechanism. In support, genetic variants may affect promoter shape for ubiquitously expressed genes in flies with limited effect on promoter expression (Schor et al., 2017). Changes in promoter shape in flies thus likely recapitulates the plasticity in TSS usage across human LCLs, despite apparent differences in core promoter elements, promoter nucleotide content, and regulatory features between flies and humans.

Notably, many of the promoter features we, and others (Sigalova et al., 2020), have identified to be predictive of promoter variability, including the presence or absence of CpG islands and TATA boxes, have previously been linked with different levels of transcriptional noise as inferred from single-cell experiments (Faure et al., 2017; Morgan and Marioni, 2018). This suggests that variability in promoter activity across individuals partly reflects the stochasticity in gene expression across cells. Given that the underlying sources of variation are different, e.g., genetic and environmental versus stochastic, this indicates that mechanisms that contribute to the buffering of stochastic noise at a single cell level can also contribute to the attenuation of genetic and environmental variation at an individual level.

We observed less restrictive +1 nucleosome positioning across individuals at low variable promoters with flexible TSS usage. Notably, these promoters are also associated with a fuzzy +1 nucleosome positioning across cells within an individual. This indicates that positional shifts of nucleosomes in coordination with shifts in core promoter usage may be due to an inherent property of these promoters in addition to the influence by genetic variation. Our observations indicate that multiple configurations of accessible chromatin may exist for low variable promoters across single cells, which may cause stochastic TSS selection with no or only low impact on expression level. This is compatible with a high density of pyrimidine/purine (YR) dinucleotides within broad promoters (Carninci et al., 2006; Frith et al., 2007), which provide a flexibility for transcription initiation sites in the absence of strong positional signals like the TATA box (Carninci et al., 2006; FANTOM Consortium and the RIKEN PMI and CLST (DGT) et al., 2014; Müller and Tora, 2014). Genetic variants biasing such TSS selection and the preference for any open chromatin configuration may therefore cause observable shifts in TSS usage between individuals, enabled by the flexible nucleosome and TSS architecture of the promoter.

It is important to note that the regulatory programs of EBV-immortalized LCLs, like other cultured cells, have been shown to be susceptible to genotype-independent sources of variation, such as primary cellular heterogeneity and EBV’s viral cellular reprogramming (Choy et al., 2008; Ozgyin et al., 2019). While we cannot exclude that such sources may influence the measured variability of some genes, we speculate that non-genetic variation would rather dampen than elevate the associations and effect sizes we report here, in particular the decomposed promoter switches highlighted by our genotype association analysis.

Taken together, our results favor a model in which the regulation of transcriptional noise across single cells reflects specificity across cell types and dispersion across individuals with shared mechanisms conferring stochastic, genetic and environmental robustness (Fig. 6). There are several implications of this model. First, the link between low transcriptional noise and low individual variability of promoters and their associations with ubiquitous and essential genes indicate that regulatory mechanisms that ensure broad expression across cell types may enforce low variability across individuals and single cells. Second, our results indicate that encoding responsiveness or developmentally restricted expression patterns of gene promoters may require high stochasticity in expression across single cells, which in turn may disallow ubiquitous expression across cell types. Thus, it is likely that increased variability is not just reflecting the absence of regulatory mechanisms that attenuate variability but the presence of those that amplify it. Finally, given that mutational robustness through flexible TSS usage is mostly associated with low variable genes, this implies that cell-type restricted, responsive and developmental genes may be more susceptible for trait-associated genetic variants, which finds support in the literature (Finucane et al., 2015; Kundaje et al., 2015; Timshel et al., 2020).

**Figure 6:**
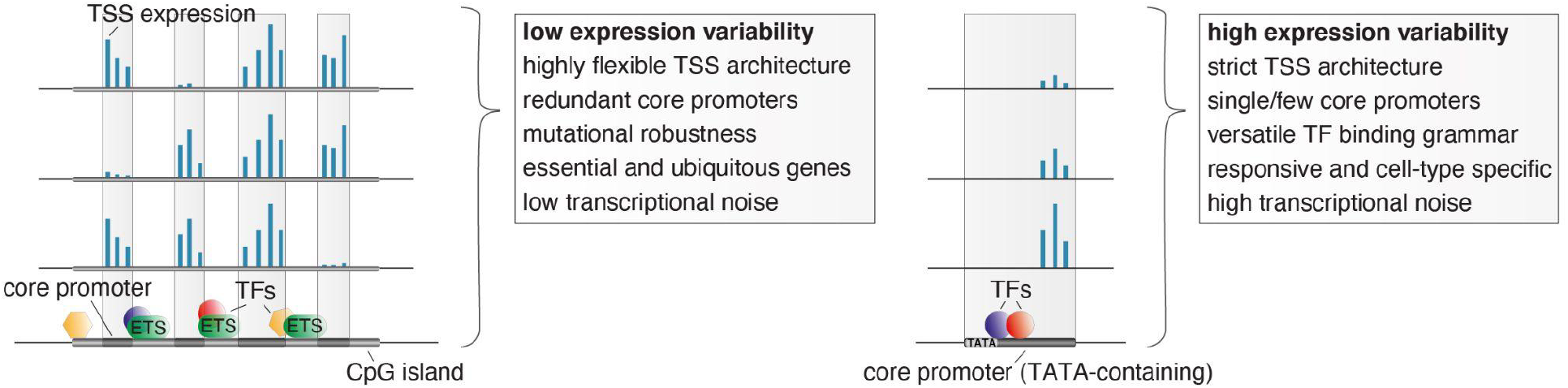
Unifying mechanisms influencing the variability in expression across individuals, the specificity in expression across cell types, and the stochasticity in expression across individual cells. Low variable promoters (left) are frequently associated with high CpG content (CpG islands), multiple binding sites of ETS factors, and a highly flexible transcription initiation architecture arising from multiple redundant core promoters (decomposed promoters) in a permissive nucleosome positioning environment. These stabilizing features along with a less complex TF binding grammar likely also act to buffer transcriptional noise across single cells and cause ubiquitous expression across cell types. The flexibility in redundant core promoter activities confers a novel layer of mutational robustness to genes. Highly variable promoters (right), on the other hand, are associated with a highly versatile TF regulatory grammar, TATA boxes, and low flexibility in TSS usage. These features likely cause, in addition to high expression variability between individuals, a responsiveness to external stimuli, cell-type restricted activity, high transcriptional noise across single cells, and less tolerance for genetic variants.

## Materials and methods

### LCL cell culturing

Epstein-Barr virus immortalized LCLs (Supplementary Table 1) were obtained from the NIGMS Human Genetic Cell Repository at Coriell Institute for Medical Research. Cells were incubated at 37°C at 5% carbon dioxide in the Roswell Park Memorial Institute (RPMI) Medium 1640 supplemented with 2mM L-glutamine and 20% of non-inactivated fetal bovine serum and antibiotics. Cell cultures were split every few days for maintenance. All 108 LCLs were grown unsynchronized for 5-7 passages and harvested when they reached >20 million cells. As these cell lines were freshly purchased, mycoplasma contamination screening and cell line authentications were not undertaken in house.

### CAGE library preparation, sequencing, and mapping

CAGE libraries were prepared in 10 batches in total as described elsewhere (Andersson et al., 2014b; Takahashi et al., 2012) from 1,500 ng total RNA from each LCL. 23 libraries (Supplementary Table 1) underwent a second round of size selection (Invitrogen E-Gel) to remove excessive primer dimers. The libraries were quality checked using an Agilent 2100 Bioanalyzer system with a RNA pico chips kit (Agilent) and quantified using DNA 1000 chips kit (Agilent). Pooled libraries (Supplementary Table 1) were sequenced with spiked-in PhiX on an Illumina HiSeq 2500 machine single-end for 50 cycles using v4 sequencing chemistry (Illumina Inc.) and a custom sequencing primer (Takahashi et al., 2012). Libraries were split by barcode and reads were trimmed to remove linker sequences and filtered for a minimum sequence quality of Q30 in 50% of base pairs using the FASTX-Toolkit. rRNA reads matching subsequences of the human ribosomal DNA complete repeating unit (U13369.1) were removed using rRNAdust (version 1.06) (FANTOM Consortium and the RIKEN PMI and CLST (DGT) et al., 2014). Mapping to the human reference genome (hg38) was performed using BWA (version 0.7.15-r1140) allowing a maximum edit distance of 2. To reduce mapping bias, reads were re-mapped using the WASP pipeline (van de Geijn et al., 2015) and BWA, taking into account biallelic SNVs (Lowy-Gallego et al., 2019). Reads with a mapping quality of 20 were retained for further analyses. Sample-related information, including CAGE run batch ID and E-gel information, are provided in Supplementary Table 1.

### CAGE tag clustering, filtering and quantification

CAGE-defined transcription start sites (CTSSs) were identified from 5’ ends of mapped CAGE reads for each strand separately. The expression of CTSSs for each LCL was quantified from the number of CAGE reads sharing 5’ ends using CAGEfightR (version 1.10) (Thodberg et al., 2019). To identify broad promoters that could potentially encompass multiple alternative core promoters (decomposed promoters), we performed lenient positional clustering of CTSSs (tag clustering) (Carninci et al., 2006), grouping CTSSs on the same strand within 60 bp of each other. To exclude rare promoters within the panel, tag clustering was performed on CTSSs with at least 1 CAGE read in at least 5 LCLs. The expression of each tag cluster (CAGE-inferred promoter) in each individual LCL was quantified by aggregating the expression of all CTSSs falling within the defined tag cluster region. To allow capture of flexible TSS usage within promoters across the panel, no support filtering was performed at CTSS level for expression quantification. Expression levels were converted to tags per million (TPM), by normalizing the expression count of each tag cluster in each library as a fraction of its number of mapped CAGE reads, scaled by 10^6^. Tag clusters were filtered to be proximal to GENCODE-annotated TSSs (hg38, version 29, within 1000bp upstream) and to have at least 10 read counts in more than 10 LCLs. The resulting 29,001 gene-associated CAGE-inferred promoters were later decomposed by local maxima decomposition to split multi-modal tag clusters (https://github.com/anderssonlab/CAGEfightR_extensions, version 0.1.1). First, for each CAGE-inferred promoter, local maxima of within-promoter CTSSs with the highest pooled expression separated by at least 20 bp were identified. Second, decomposition was performed for each local maxima separately in decreasing order of pooled expression level. For each local maxima, the fraction between the pooled expression of each CTSS to that of the local maxima was calculated. All CTSSs associated with at least 10% of the local maxima signal that were not gapped by more than 10 bp with CTSS expression less than this value were retained in a new decomposed promoter. For smoothing purposes, neighboring non-zero CTSSs within 1 bp distance of CTSSs fulfilling the fraction criterion were also included. Subsequently, decomposed promoters were merged if positioned within 1 bp from each other.

### Geuvadis YRI RNA-seq data analysis

Gene expression data quantified in the recount2 project (Collado-Torres et al., 2017) using Geuvadis YRI RNA-seq data (Lappalainen et al., 2013) was downloaded using the recount R package. Only genes with more than 1 transcript per million in at least 10% of YRI samples were considered for expression variability calculation.

### GM12878 scRNA-seq data analysis

GM12878 10X Genomics scRNA-seq data (Osorio et al., 2019) was downloaded from Gene Expression Omnibus (GSE126321) and processed using Seurat (version 4.0.3) (Hao et al., 2021). Cells with a proportion of mitochondrial reads lower than 10% and a sequencing depth deviating less than 2.5 times the standard deviation from the average sequencing depth across cells were considered. The expression of genes with read counts observed in at least 10 cells were normalized using scran (version 1.18.7) (Lun et al., 2016) and retained for expression variability calculation.

### Measuring expression variability across individuals

The raw dispersion of each CAGE tag cluster was calculated using the squared coefficient of variation (CV^2^) of TPM-normalized promoter (or decomposed promoter) expression across the LCL panel and subsequently log_10_-transformed. Adjustment of the mean expression-dispersion relationship was performed by subtracting the expected log_10_-transformed dispersion for each promoter according to its expression level, using a running median (width 50, step size 25) of raw dispersions (log_10_ CV^2^) ordered by mean expression level (TPM) across the panel, as described elsewhere (Kolodziejczyk et al., 2015; Newman et al., 2006). The same strategy was used to calculate the adjusted dispersion of gene expression from RNA-seq and scRNA-seq data. Promoters were grouped by variability according to the quartiles of expression-adjusted dispersions (≤Q1, (Q1, Q2], (Q2, Q3], >Q3).

### Neural network model, training and hyperparameter tuning

A simple neural network architecture was designed to learn to predict low and high variability from DNA sequence. The neural network model uses as input one-hot-encoded DNA sequences (A = [1,0,0,0], C = [0,1,0,0], G = [0,0,1,0], T = [0,0,0,1]) from the human reference genome (hg38) to predict low and highly variable promoter activity as output. Although CAGE-inferred promoters varied in width, we made use of fixed-length 600 bp sequences for each promoter centered on its pooled CAGE summit CTSS. 600 bp was used to make sure that sequences influencing promoter variability contained within regions that could cover a central open chromatin site (150-300bp) as well as within flanking nucleosomal DNA (150-200bp) were captured, where also most of the expression output of a promoter originate (FANTOM Consortium and the RIKEN PMI and CLST (DGT) et al., 2014).

The model (Supplementary Fig. 2A) consists of one convolutional layer with 128 hidden units and a kernel size of 10, followed by global average pooling and two dense layers with 128 and 2 nodes, respectively. Batch normalization and dropout (0.1) were applied after each layer. The ReLU activation function (Agarap, 2019) was used in all layers except the final layer, in which a sigmoid activation function was used to predict the variability class (low or high adjusted dispersion).

Promoter sequences from chromosomes 2 and 3 were used as the test set and those from the remaining chromosomes were used for training and hyperparameter tuning with a 5-fold cross-validation. Hyperparameters were manually adjusted to yield the best performance on the validation set. The neural network model was implemented and trained in Keras (version 2.3.0, https://github.com/fchollet/keras) with TensorFlow backend (version 1.14) (Abadi et al., 2016) using the Adam optimizer (Kingma and Ba, 2017) with a learning rate of 0.0001, batch size of 64, and early stopping with the patience of 15 epochs.

We initially used the first and third quartiles (Q1 and Q3) to distinguish low variable promoters (≤Q1) from highly variable promoters (>Q3), corresponding to an adjusted log_10_-transformed CV^2^ of −0.1490 and 0.1922, respectively. To reduce false positives, we slightly adjusted the thresholds for low and highly variable promoters to −0.20 and 0.25, respectively. The final training and test sets for the neural network model together consisted of 5,054 low variable and 5,683 highly variable promoters. To ensure consistency, the same thresholds were used for training and testing with Random Forest and decision tree classifiers (see below).

### Motif discovery using DeepLIFT and TF–MoDISco

To interpret the neural network model we used DeepLIFT (Shrikumar et al., 2019), a feature attribution method, to compute importance scores for each nucleotide in the 600bp input sequences for low and highly variable promoters. DeepLIFT relies on backpropagation of the contributions of all neurons in the neural network to the input features, nucleotides, and was used to estimate the importance of each position and nucleotide in the input sequences to predict high and low variability. The resulting importance scores were supplied to TF-MoDISco (Transcription Factor Motif Discovery from Importance Scores) (Shrikumar et al., 2020) to identify DNA stretches (seqlets) with high importance for the predictions. DeepLIFT and TF-MoDISco were run independently on the input sequences for low variable and highly variable promoters. TF-MoDISco identified 18,035 seqlets for low variable promoters and 21,942 seqlets for highly variable promoters by using the importance scores from DeepLIFT over a width of 15 bp with a flank size of 5 bp and a FDR threshold of 0.05. The seqlets identified were merged in 41 and 47 metaclusters for low and highly variable promoters, respectively.

We used Tomtom (MEME package 5.1.1) (Gupta et al., 2007) to match the resulting metaclusters to known TF motifs (in MEME format) from the JASPAR database (release 2020, hg38) (Fornes et al., 2020). We compared each non-redundant JASPAR vertebrate frequency matrix with the metaclusters using Tomtom based on the Sandelin and Wasserman distance (Sandelin and Wasserman, 2004). Matches were considered those with a minimum overlap between query and target of 5 nucleotides and a p-value < 0.05.

### Random forest, Boruta and SHAP analysis

To identify broad-scale trends of high CpG content, we calculated CpG observed/expected ratio in windows +/− 500bp around the pooled summit CTSS of each promoter. Calculated CpG ratios revealed a bimodal distribution that informed on thresholding high CpG content promoters as those with CpG observed/expected ratio >0.5 (Supplementary Fig. 3A).

Predicted transcription factor binding sites for 746 TFs with scores of 500 or greater (P < 10^−5^) (hg38) were obtained from JASPAR (release 2020, hg38) (Fornes et al., 2020) and for each TF, presence/absence was obtained by overlapping predicted TF binding sites with promoters considered in the modeling. Together, the CpG content status and the presence/absence of predicted TF binding sites were used as features for predicting high and low variability using Random Forest (Pedregosa et al., 2011).

Similarly to the neural network model, promoters from chromosomes 2 and 3 were only used as the test set. The remaining promoters were used for training and hyperparameter tuning with 5 fold cross-validation. The Random Forest model was implemented and trained in Scikit-learn (version 2.3.0) with 500 trees, a maximum depth of trees of 10, 50 samples split per node, and 50 samples to be at a leaf node. The remaining hyperparameters were kept with default values.

Instead of selecting features directly from the Random Forest model, the BorutaShap package (Keany, 2020) was used for feature selection. The main advantage of using the Boruta approach is that the features compete with their randomized version (or shadow feature) and not with themselves. Therefore, a feature is considered relevant only if its score is higher than the best randomized feature. In this way, from the 746 original TF features, only 125 features were kept. The features were selected using only promoters from the training set. Finally, the SHAP library (Lundberg and Lee, 2017) was used to explain the importance of the 125 selected features for the two promoter classes. SHAP calculates Shapley values, a game theoretic approach for optimal credit allocation during cooperation, which can be used to estimate the marginal contribution of each feature to a model’s predictions.

### Decision tree baseline model

To evaluate the contribution of TF binding site presence for predicting promoter variability, we trained a baseline model based on CpG content status and TATA-box presence only. CpG content status (CpG observed/expected ratio > 0.5) and the presence/absence of predicted TBP binding sites were used as features for predicting high and low variability using a decision tree classifier. The decision tree model was implemented and trained in Scikit-learn (version 2.3.0) using default parameters. Training and test data were defined as for the CNN and Random Forest models.

### Tissue-, cell-type specificity and gene annotations

RNA-seq gene expression values across tissues were obtained from the GTEx consortium (GTEx Consortium, 2017). Promoters were considered expressed in tissues in which their corresponding gene had ≥ 5 RPKM average expression across donors.

CAGE gene expression values across cell types were obtained from the FANTOM5 project (FANTOM Consortium and the RIKEN PMI and CLST (DGT) et al., 2014). The average normalized (tags per million, TPM) expression per gene was calculated across samples associated with the same cell type facet (grouping of CAGE libraries according to Cell Ontology annotation of samples), according to (Andersson et al., 2014a), and a gene was considered expressed in a cell type facet if the average expression was ≥ 5 TPM.

Gene lists for FDA approved drug-targets (Wishart et al., 2018), essential genes (Hart et al., 2017) and GWAS hits (MacArthur et al., 2017) were downloaded from the MacArthur Lab Repository (https://github.com/macarthur-lab/gene_lists).

### GM12878 cell culturing, TNF-α stimulation and differential expression analysis

GM12878 cells were obtained from the NHGRI Sample Repository for Human Genetic Research at Coriell Institute for Medical Research. Unstimulated GM12878 cells and those stimulated with 25ng/ml TNF-α for 6 hours were harvested with four replicates for each condition. Cell culturing, CAGE library preparation and mapping were done as described above for the LCL panel. CAGE reads supporting each of the final filtered promoters identified in the LCL panel were counted for each replicate using CAGEfightR (version 1.10) (Thodberg et al., 2019). Differential expression analysis of the aggregated CTSS counts was performed using standard library size adjustment and a generalized linear model with DESeq2 (version 1.30.1) (Love et al., 2014). Promoters with FDR-adjusted p-value ≤ 0.05 were considered to be differentially expressed.

### Correlation analysis of decomposed promoter expression

To test if decomposed promoters could act independently of each other, we calculated Pearson correlation coefficients of LCL expression between pairs of decomposed promoters originating from the same promoter. We focused on promoters with at least 2 decomposed promoters significantly contributing to the overall expression of the promoter. Specifically, we considered decomposed promoters whose expression accounted for at least 5% of the overall promoter expression in at least half of all LCLs, resulting in 37,663 decomposed promoters of 14,889 promoters. To avoid potential bias introduced from a variable number of decomposed promoters per promoter, we considered the lowest correlation across decomposed promoter pairs within a promoter.

#### Nucleosome positioning analysis

Micrococcal nuclease-digested nucleosome sequencing (MNase-seq) data from 7 EBV-immortalized LCLs (GM18507, GM18508, GM18516, GM18522, GM19193, GM19238, GM19239) were obtained from GEO (GSE36979) (Gaffney et al., 2012).

The locations and fuzziness scores of nucleosomes were called with DANPOS2 (version 2.2.2) (Chen et al., 2013) using the dpos command on each LCL separately. +1 nucleosomes were defined as the closest downstream nucleosome of the dominant TSS position (derived from pooled CAGE data) of each CAGE-inferred promoter.

Positional cross correlations were calculated between CAGE TSSs and 5’ ends of MNase-seq reads on the same strand at dominant TSS positions of CAGE-inferred promoters ±500 bp (maximum lag 250) to identify their most likely separation. Cross-correlation analysis was performed on either pooled CAGE data (across all 108 LCLs) versus pooled MNase-seq data (across all 7 LCLs) or using only CAGE and MNase data from one LCL (GM18516). Finally, for both analyses, a weight for each promoter was calculated from the geometric mean of aggregated MNase and CAGE signals. This was used to calculate a weighted average of the cross correlations across considered promoters.

### Mapping QTLs

We tested both the association between the genotypes of common genetic variants (MAF ≥ 10%) and the expression of promoters (promoter eQTL, prQTLs) and decomposed promoters (decomposed promoter eQTL, dprQTLs), as well as their association with the contribution of to the overall expression of the encompassing (non-decomposed) promoter (fraction eQTL, frQTL). prQTLs, dprQTLs, and frQTLs were mapped using the MatrixEQTL R package (version 2.3) (Shabalin, 2012). We controlled for genetic population stratification and library preparation batches (Supplementary Table 1) by including these as covariates. In addition, we included the first 5 principal components derived from normalized promoter expression values (TPM) as covariates for prQTLs.

For prQTL detection, all 29,001 promoters were tested using TPM-normalized expression values. For frQTLs, we calculated the fractional contribution of each decomposed promoter to the expression of its original promoter. To focus the dprQTL and frQTL analyses on relevant shifts in TSS usage, we considered only decomposed promoters whose expression accounted for at least 5% of the overall promoter expression in at least half of all LCLs and promoters with at least 2 such decomposed promoters, resulting in 37,663 decomposed promoters of 14,889 promoters.

For each promoter, we tested common (minor allele frequency ≥10%) biallelic SNVs (Lowy-Gallego et al., 2019) at a maximum distance of 25kb from the CTSS with maximum pooled CAGE signal within each promoter for association with changes in promoter expression levels or decomposed promoter contribution. Resulting p-values were FDR-adjusted according to the total number of promoters or decomposed promoters tested genome-wide within the MatrixEQTL R package. prQTLs, dprQTLs, and frQTLs with FDR ≤ 5% were retained. A promoter was associated with a dprQTL or frQTL if at least one of its decomposed promoters was associated with a dprQTL or frQTL at FDR < 5%.

## Code availability

Code for data analysis performed in this study is publicly available on GitHub: https://github.com/anderssonlab/Einarsson_et_al_2022/.

## Data availability

CAGE data were deposited into the Gene Expression Omnibus (GEO) database under accession number GSE188131 (https://www.ncbi.nlm.nih.gov/geo/query/acc.cgi?acc=GSE188131). GM12878 scRNA-seq data were retrieved from GEO (accession number GSE126321). Processed Geuvadis RNA-seq gene expression data were retrieved from recount2 (Collado-Torres et al., 2017) (accession number ERP001942). Processed GTEx RNA-seq gene expression data were retrieved from the GTEx portal (https://www.gtexportal.org/home/datasets, version-8). Predicted transcription factor binding sites for 746 TFs were obtained from JASPAR 2020 (http://expdata.cmmt.ubc.ca/JASPAR/downloads/UCSC_tracks/2020/hg38/).

## Acknowledgements

We thank members of the Andersson lab for rewarding discussions. Sequencing was performed by the SNP&SEQ Technology Platform in Uppsala, part of the National Genomics Infrastructure (NGI) Sweden and Science for Life Laboratory. The SNP&SEQ Platform is also supported by the Swedish Research Council and the Knut and Alice Wallenberg Foundation.

This work was supported by funding from the Danish Council for Independent Research [grant 6108-00038], the European Research Council (ERC) under the European Union’s Horizon 2020 research and innovation programme [grant 638173], and the Novo Nordisk Foundation [grants NNF18OC0052570, NNF20OC0059796].

## Author contributions

H.E. and R.A. conceived the project; H.E. led the data analysis with support from N.A., S.R., and R.A.; M.S. performed machine learning; C.V. performed CAGE experiments with contributions from J.B.L.; R.A. supervised the project; H.E. and R.A. wrote the manuscript; all authors reviewed the final manuscript.

## Notes

### Competing Interest Statement

The authors have declared no competing interest.

### Summary of Updates

We have revised the manuscript with additional analyses and discussions of alternative interpretations, to address reviewers' questions and comments regarding nucleosome positioning, initiator strength, and decomposed promoter eQTLs. We have further revised parts of the text for clearer and a more cautious language where appropriate.

