## Supplementary material for "Promoter sequence and architecture determine expression variability and confer robustness to genetic variants"

**Contents**

Supplementary Figures 1-10
Legends for Supplementary Tables 1-9

### Supplementary Figures

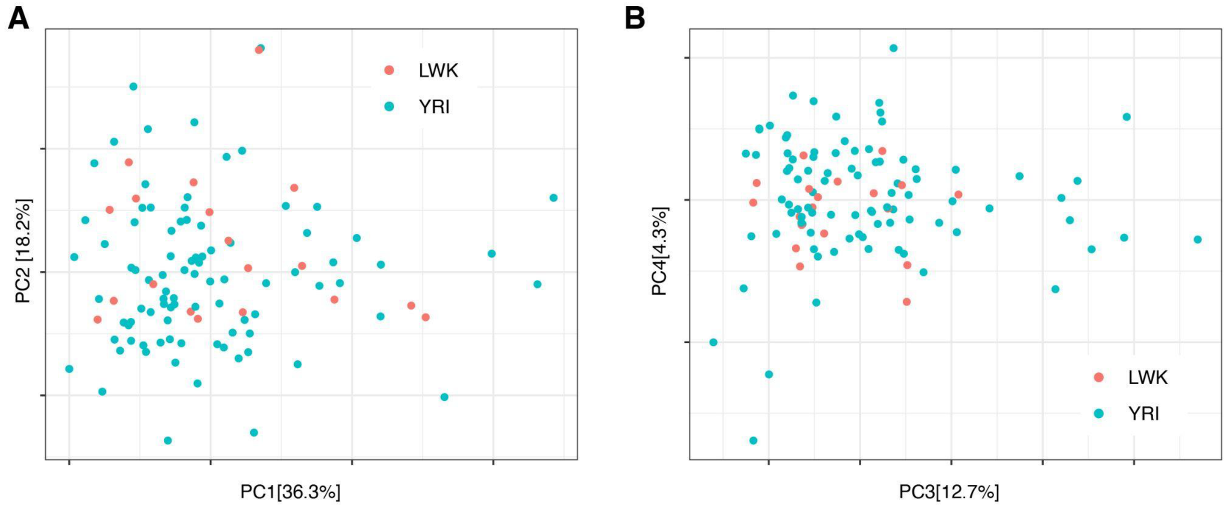


**Supplementary Figure 1: PCA plot of promoter expression (CAGE) across the LCL panel.** 1st and 2nd (A), and 3rd and 4th (B) principal components (PCs), colored according to population (YRI and LWK). PCA was performed using TPM-normalized expression for all 29,001 considered promoters. Percentage of variation accounted for by each principal component is shown in brackets with the axis label.

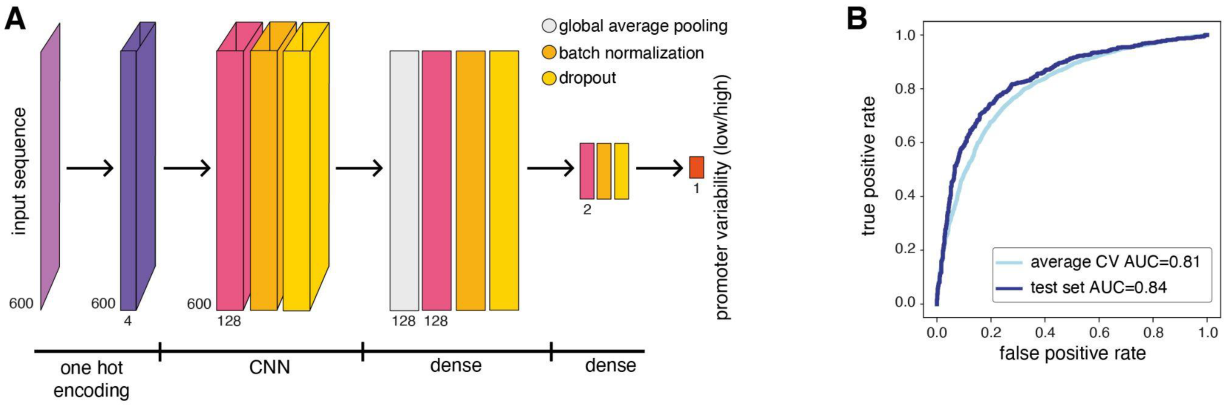


**Supplementary Figure 2: Neural network model and performance. A:** Neural network architecture used for learning promoter variability from promoter sequence. The architecture is composed of one convolutional layer with 128 hidden units, followed by global average pooling and two dense layers with 128 and 2 nodes, respectively. **B:** Receiver-operating curves (ROC) for average cross validation (light blue, AUC=0.81) and the test set (dark blue, AUC=0.84).

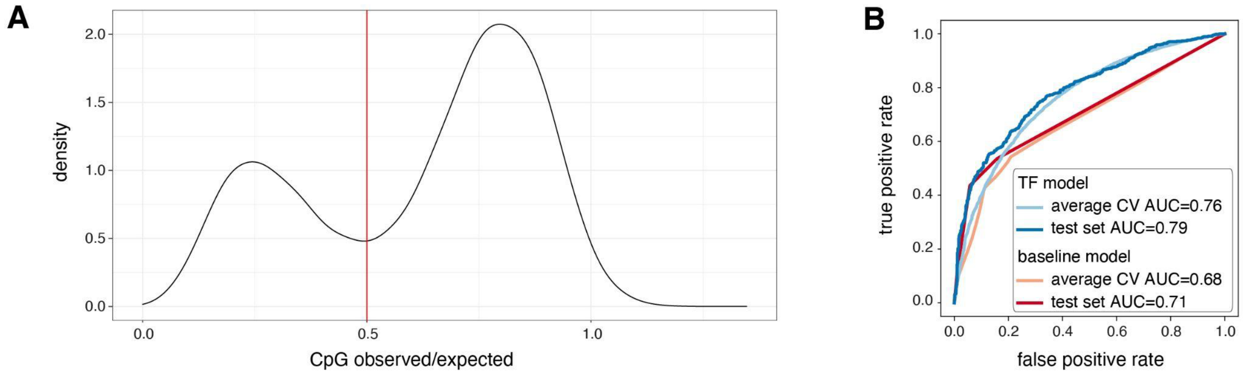


**Supplementary Figure 3: Random forest features and performance. A:** Observed / expected CpG ratio calculated in windows covering +/- 500 bp around the CAGE summit position of considered promoters. Red vertical line marks the threshold (0.5) between low and high CpG content. **B:** Random forest model (TF model) receiver-operating curves (ROC) for average cross validation (light blue, AUC=0.76) and the test set (dark blue, AUC=0.79). Shown are also the ROC for a decision tree (baseline) model based on CpG content and TBP binding sites alone (average cross validation AUC=0.68, orange; test set AUC=0.71, red).

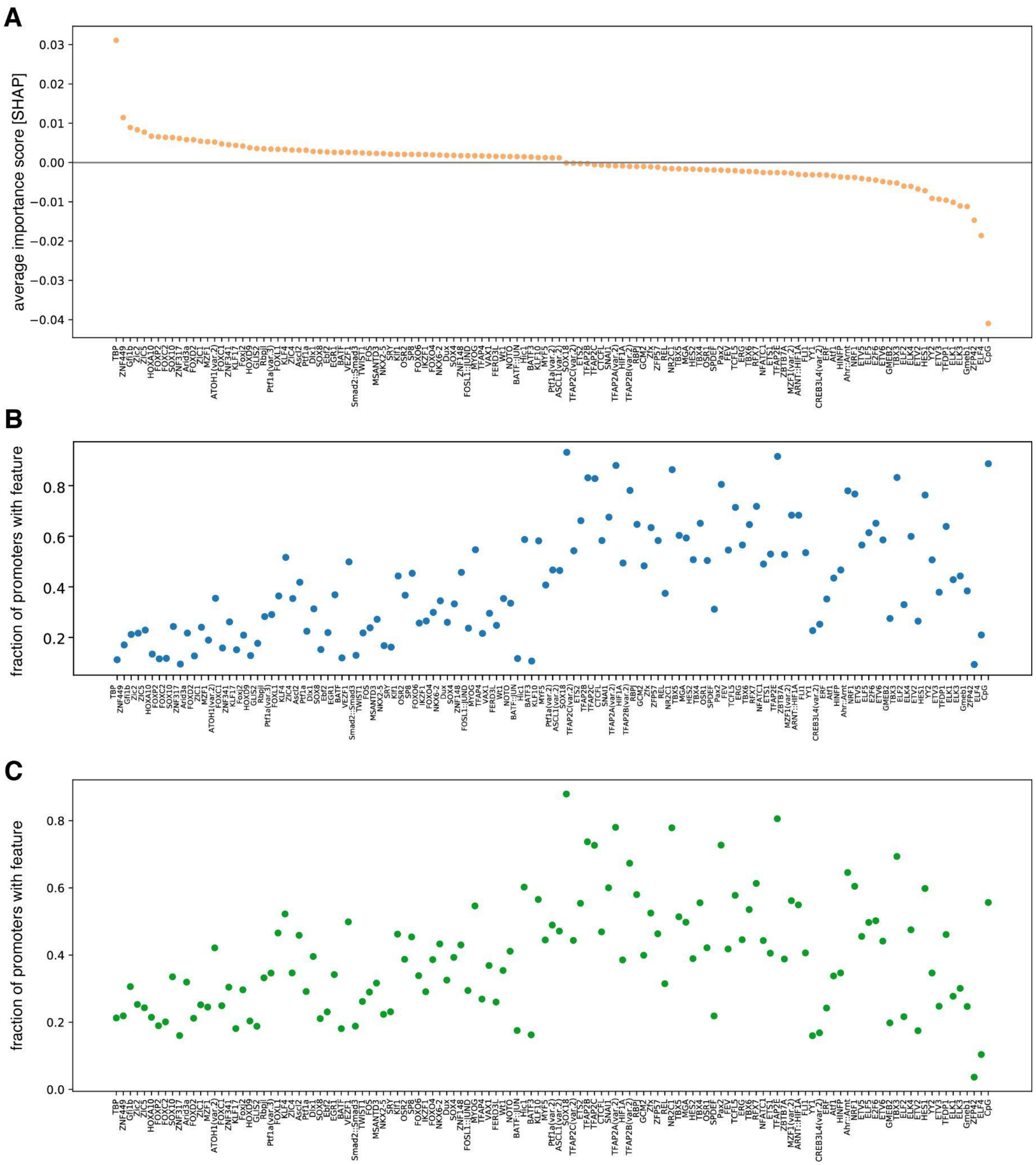


**Supplementary Figure 4: Full panel of features found to be predictive of promoter variability. A:** Average contribution (SHAP values) of each of the 125 TFs identified as important for predicting promoter variability. TFs are ordered by their average contribution to the prediction of highly variable promoters. **B-C:** The frequency of predicted TF binding sites (presence/absence) in low variable (B) and highly variable (C) promoters. TFs follow the same order as in panel A.

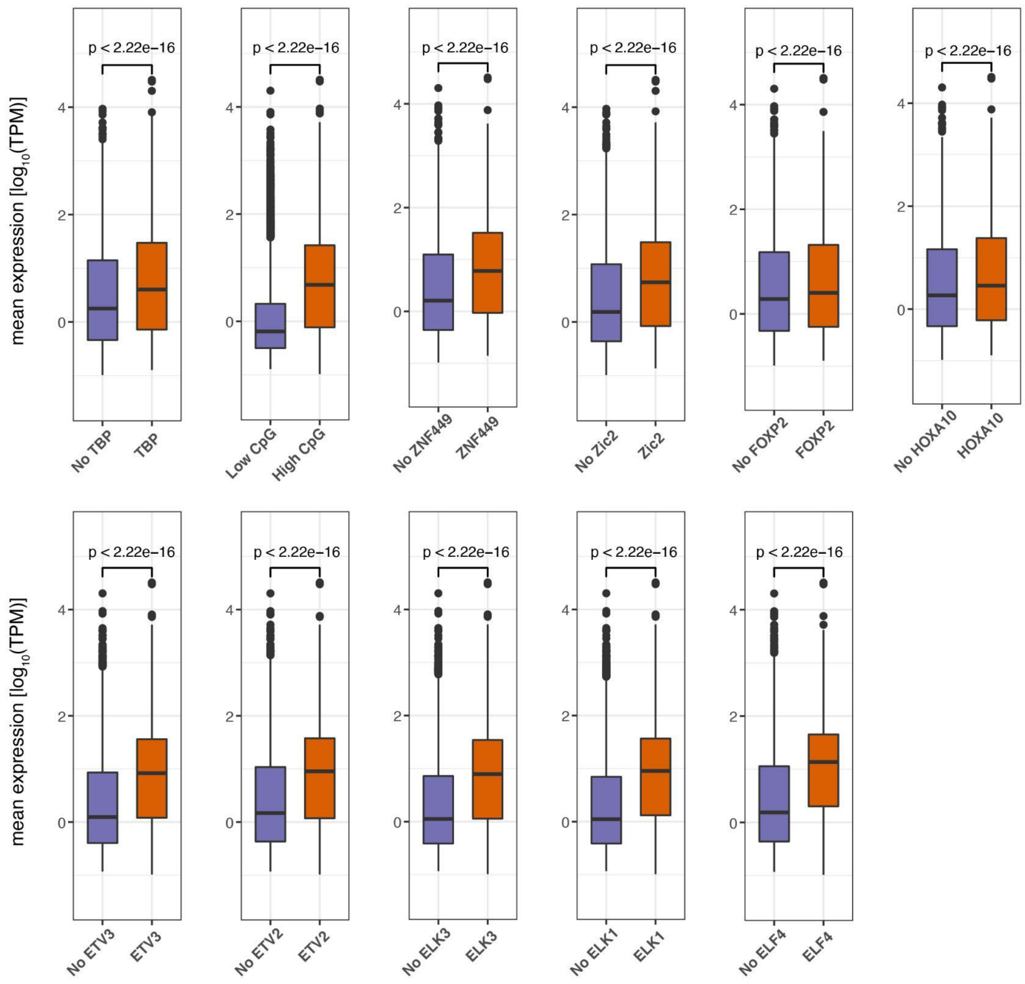


**Supplementary Figure 5: Association between TF binding sites and promoter expression level.** Box-and-whisker plots displaying the difference in TPM normalized promoter expression between in the absence (blue) or presence (orange) of TF binding sites. For all box-and-whisker plots, central band: median; boundaries: first and third quartiles; whiskers: +/- 1.5 IQR. P-values were determined using the Wilcoxon rank-sum test (***: p<0.05).

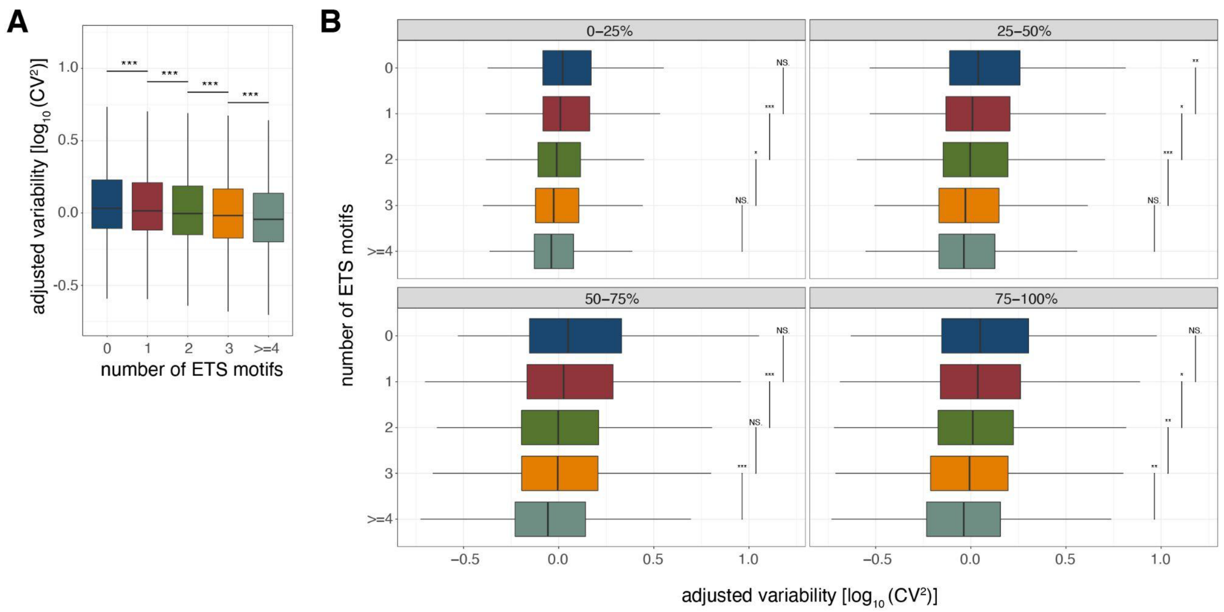


**Supplementary Figure 6: Association between the number of ETS binding sites and promoter variability. A:** Variability of promoters grouped by their number of predicted non-overlapping ETS binding sites. **B.** Variability of promoters grouped by their number of predicted non-overlapping ETS binding sites split based on promoter expression level quartiles. For all box-and-whisker plots, central band: median; boundaries: first and third quartiles; whiskers: +/- 1.5 IQR. In both panels, outliers were not plotted. P-values were determined using the Wilcoxon rank-sum test (***: $p<0.05$).

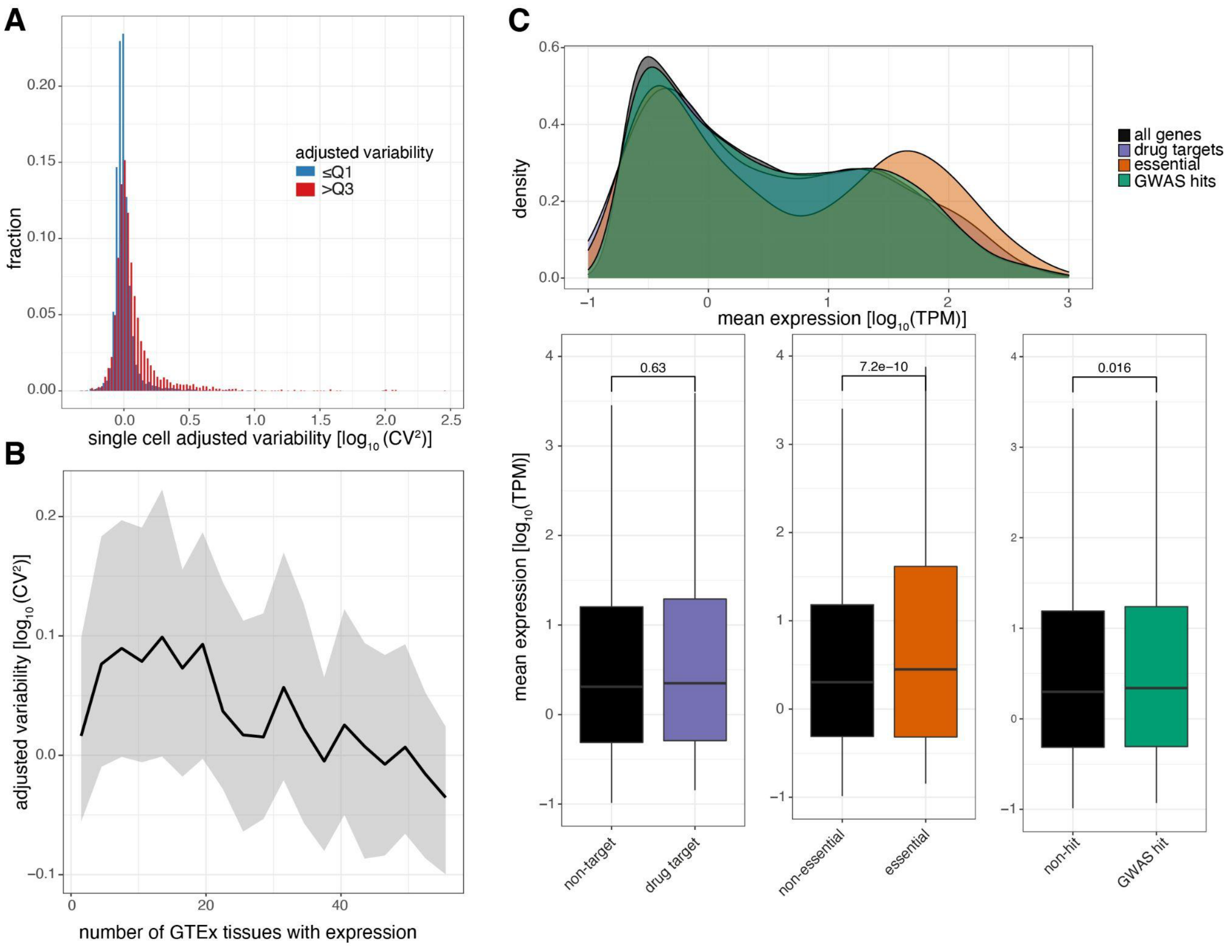


**Supplementary Figure 7: Levels of promoter variability are reflective of distinct biological processes. A:** Distribution of single cell adjusted variability [$\log_{10}(CV^2)$] of genes for low variable promoters (blue) and highly variable promoters (red). **B:** Median promoter variability (line) and interquartile range (shading), as a function of the number of GTEx tissues the associated gene is expressed in (median tissue expression >5 RPKM). **C:** Distribution of promoter expression associated with drug-targets (purple), essential (orange), or GWAS hits (green) genes, compared to all promoters (black). Top: density plot of promoter expression per gene category. Bottom: Box-and-whisker plots of promoter expression split by each category of genes. P-values were determined using the Wilcoxon rank-sum test. For all box-and-whisker plots, central band: median; boundaries: first and third quartiles; whiskers: +/- 1.5 IQR.

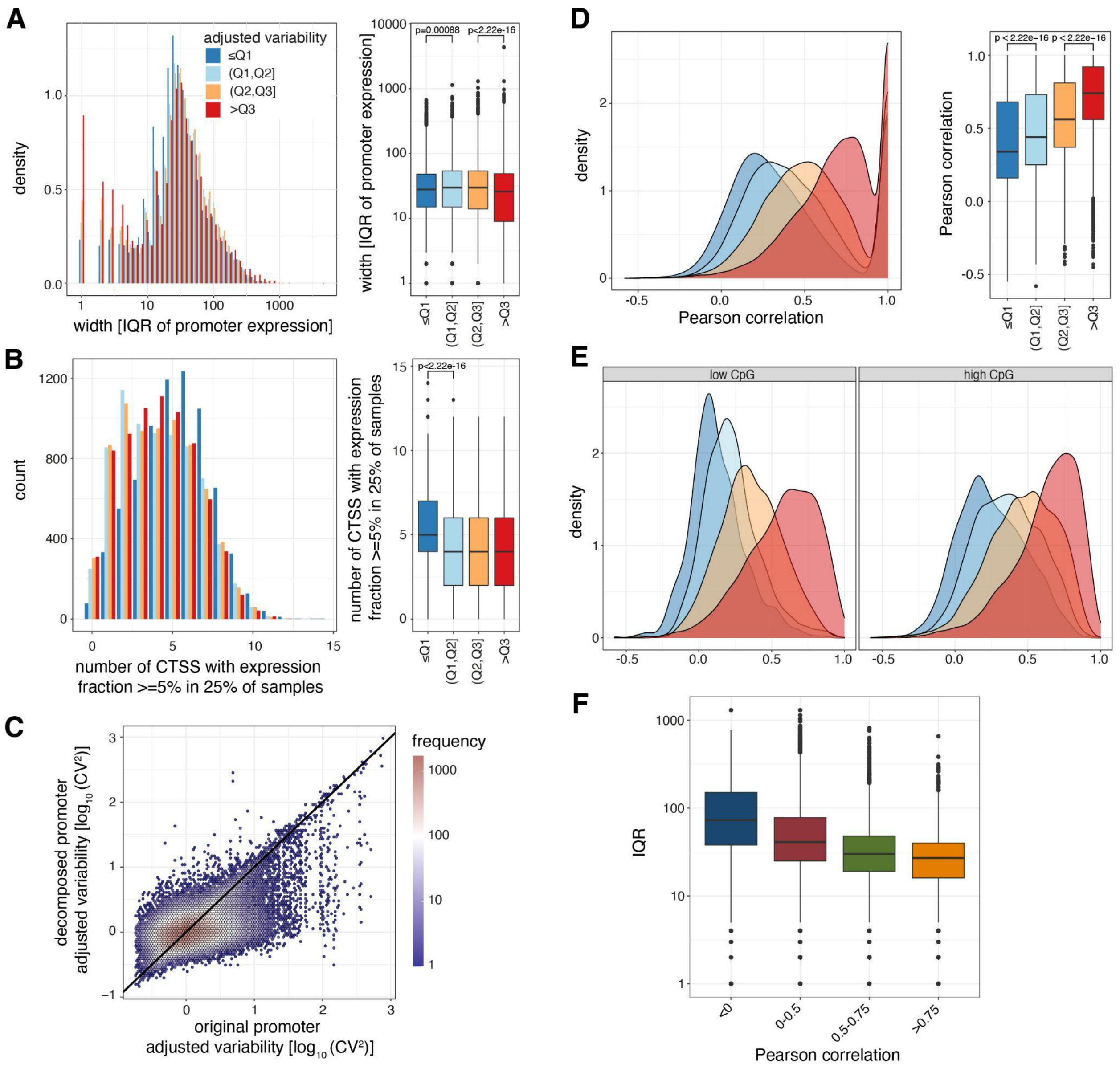


**Supplementary Figure 8: Low variable promoters tend to be composed of multiple clusters of TSSs rather than broader TSS signatures. A-B:** Promoter shape metrics for promoters split by variability quartiles. The left subpanels display the distribution of IQRs (widths containing the 25th to 75th percentiles of contained CAGE signal) (A) and the number of TSSs with expression fraction >=5% in 25% of samples (B). The right subpanels display box-and-whisker plots of the differences in these metrics across promoters split by variability quartiles (central band: median; boundaries: first and third quartiles; whiskers: +/- 1.5 IQR.). **C:** The relationship between adjusted $log_{10}$-transformed $CV^2$ of the original promoter and adjusted $log_{10}$-transformed $CV^2$ of local-maxima decomposed promoters as a 2D density chart. **D:** Densities (left) and box-and-whisker plots (right) of the Pearson correlation between all possible pairs of decomposed promoters originating from the same promoter across all CAGE-inferred promoters with at least two decomposed promoters (central band: median; boundaries: first and third quartiles; whiskers: +/- 1.5 IQR.).

**E:** Densities of the lowest Pearson correlation between any pair of decomposed promoters originating from the same promoter across all CAGE-inferred promoters with at least two decomposed promoters, split based on CpG content levels. **F:** Box-and-whisker plots of the IQRs (widths containing the 25th to 75th percentiles of contained CAGE signal) split based on levels of lowest Pearson correlation between any pair of decomposed promoters originating from the same promoter (central band: median; boundaries: first and third quartiles; whiskers: +/- 1.5 IQR).

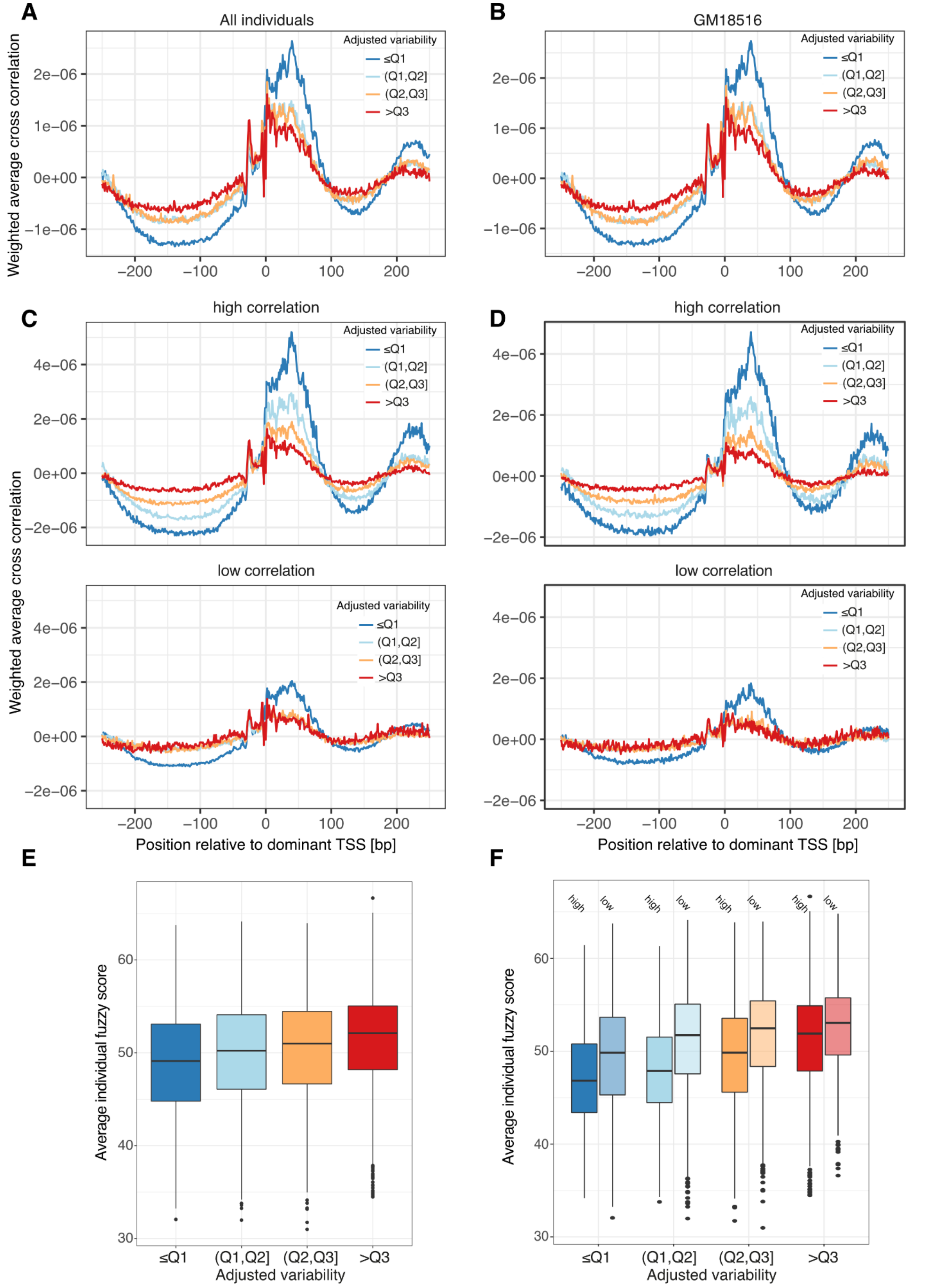

A
All individuals
B
GM18516
C
high correlation
D
high correlation
low correlation
low correlation
Adjusted variability
≤Q1
(Q1,Q2]
(Q2,Q3]
>Q3
Weighted average cross correlation
Position relative to dominant TSS [bp]
E
F
Average individual fuzzy score
Adjusted variability
high
low

**Supplementary Figure 9: Low variable promoters with highly correlated decomposed promoters tend to have a fixed +1 nucleosome position across individuals. A-D:** Weighted average cross correlation of MNase-seq nucleosome occupancy profiles relative to promoter CAGE summit positions, split by variability quartiles, for pooled CAGE data (across all 108 LCLs) versus pooled MNase-seq data (across 7 LCLs, panels: A, C) or using only CAGE and MNase data from one LCL (GM18516, panels: B, D). Weighted average cross correlations were calculated either across all promoters (A,B) or separately for promoter groups split based on their minimum correlation between contained decomposed promoter pairs (high Pearson's correlation >0.39: top panel, low Pearson's correlation <=0.39: bottom panel) (C,D). **E-F:** Average nucleosome fuzzy score across all 7 LCLs, split by variability quartiles (E) and minimum correlation between contained decomposed promoter pairs (high Pearson's correlation >0.39: solid colors, low Pearson's correlation <=0.39: faded colors) (F).

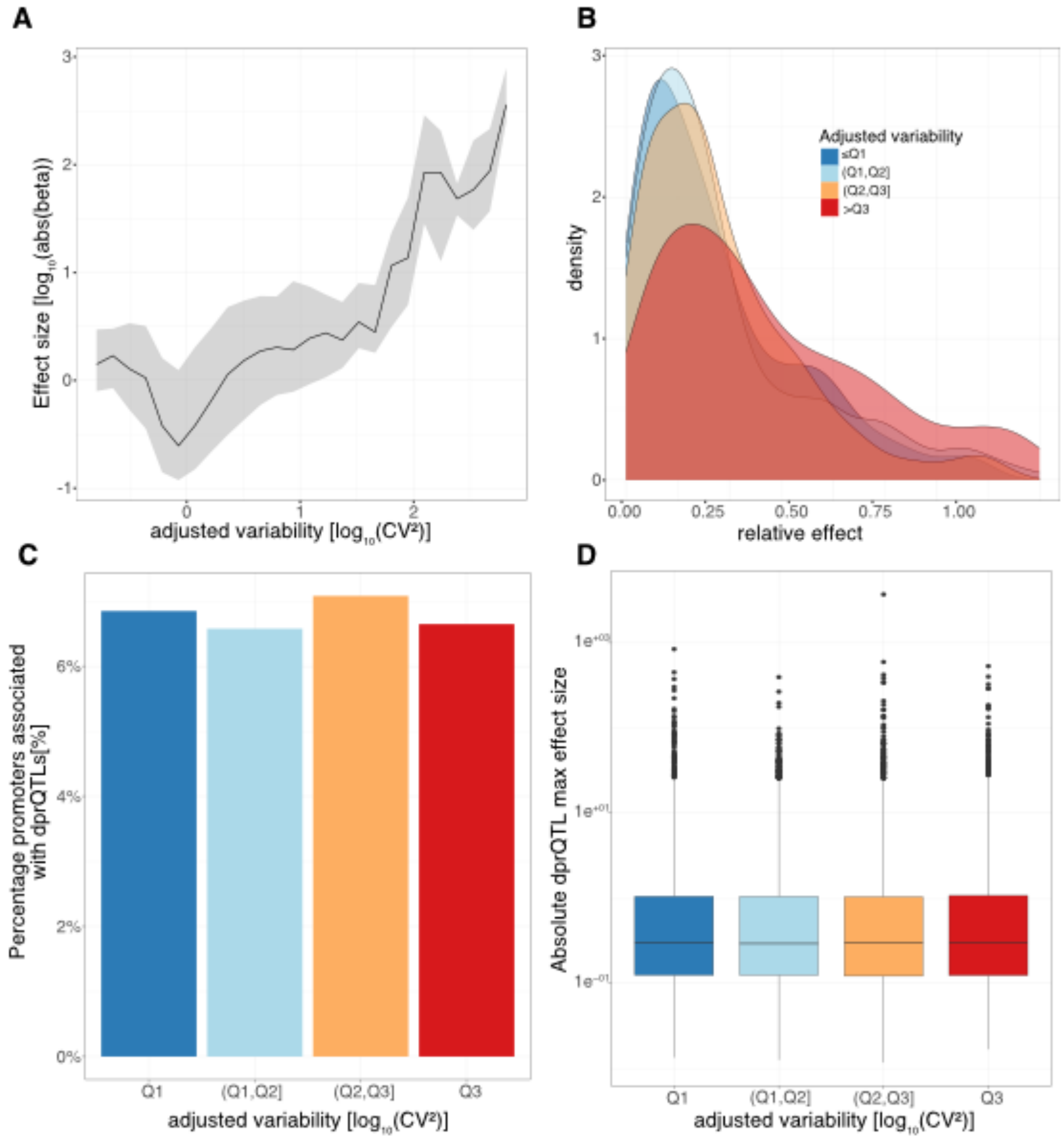


**Supplementary Figure 10: Decomposed promoters can be affected differently by proximal genetic variants compared to their encompassing promoters. A:** Median effect size [log10(abs(beta))] for the most significant prQTL for each promoter (line) and interquartile range (shading), as a function of adjusted promoter variability. **B:** Density plot of the relative effect sizes for the most significant prQTL of each promoter with FDR<=5% split by RNA-seq-derived variability quartiles. C: Percentage of promoters containing at least 1 decomposed promoter associated with a dprQTL (FDR<0.05), split by promoter variability quartiles. **D:** Maximum effect size [abs(beta)] of dprQTL each decomposed promoter, split by encompassing promoter variability quartiles.

### Supplementary Tables

**Supplementary Table 1: LCL sample and CAGE library information.**

[tab-delimited]

Row names: Cell line IDs

Columns:

- Sex: The sex of the individual that the cell line was derived from (Male/Female)
- Population: Population origin of the individual the cell line was derived from (YRI/LWK)
- E_Gel: Indicates whether samples underwent a second round of size selection (yes) or not (no)
- CageRun_ID: The CAGE library batch the sample was prepared in, with comma separating IDs if sample was included in multiple runs
- SeqPool_ID: The sequence pool IDs the sample was included in, with comma separating IDs if sample was included in multiple pools
- SeqRun_ID: The sequence run IDs the sample was included in, with comma separating IDs if sample was included in multiple sequencing runs
- Total_reads: Total reads sequenced
- Mapped_reads: Total mapped reads

**Supplementary Table 2: CAGE-inferred promoters associated with GENCODE-annotated TSSs.**

[tab-delimited]

Row names: genomic coordinates of promoters provided as chromosome:start-end;strand

Columns:

- geneID: Ensembl ID of the associated gene
- median: median TPM-normalized tag cluster expression across the LCL panel
- CV: coefficient of variation of TPM-normalized tag cluster expression across the LCL panel
- mean: mean TPM-normalized tag cluster expression across the LCL panel
- log10_CV2: $\log_{10}$-transformed squared coefficient of variation ($CV^2$) of TPM-normalized tag cluster expression across the LCL panel
- adjusted_log10_CV2: $\log_{10}$-transformed squared coefficient of variation ($CV^2$) after adjustment of the mean expression-dispersion relationship.
- adjusted_quartile: adjusted variability split by quartiles (<Q1: 0-25%, (Q1,Q2]: 25-50%, (Q2,Q3]: 50-75%, >Q3: 75-100%)

**Supplementary Table 3: Gene level expression characterization across GM12878 single cells.**

[tab-delimited]

Row names: Ensembl IDs

Columns:

- median: median scran-normalized read counts across GM12878 single cells
- CV: coefficient of variation of scran-normalized read counts across GM12878 single cells
- mean: mean scran-normalized read counts across GM12878 single cells
- log10_CV2: $\log_{10}$-transformed squared coefficient of variation ($CV^2$) of scran-normalized read counts across GM12878 single cells
- adjusted_log10_CV2: $\log_{10}$-transformed squared coefficient of variation ($CV^2$) after adjustment of the mean expression-dispersion relationship across GM12878 single cells

**Supplementary Table 4: Promoter differential expression results using DESeq2 in GM12878 after TNFα treatment.**

[tab-delimited]

Row names: genomic coordinates of promoters provided as chromosome:start-end;strand

Columns:

- baseMean: Average normalized count values
- log2FoldChange: Effect size estimate [$\log_2$ Fold Change] for 6h TNFα/untreated
- lfgSE: Standard error estimate for $\log_2$ fold change for 6h TNFα/untreated
- stat: Wald test statistics [Z-statistic]
- pvalue: Wald test p-value for 6h TNFα/untreated
- padj: Benjamin-Hochberg corrected p-value
- geneID: Ensembl ID of the associated gene
- adjusted_log10_CV2: $\log_{10}$-transformed squared coefficient of variation ($CV^2$) after adjustment of the mean expression-dispersion relationship
- adjusted_quartile: adjusted variability split by quartiles (<Q1: 0-25%, (Q1,Q2]: 25-50%, (Q2,Q3]: 50-75%, >Q3: 75-100%)

**Supplementary Table 5: Decomposed promoter expression characterization and its association with original promoter expression variability.**

[tab-delimited]

Row names: genomic coordinates of decomposed tag clusters provided as chromosome:start-end;strand

Columns:

- median: median TPM-normalized decomposed tag cluster expression across the LCL panel
- CV: coefficient of variation of TPM-normalized decomposed tag cluster expression across the LCL panel
- mean: mean TPM-normalized decomposed tag cluster expression across the LCL panel
- log10_CV2: $\log_{10}$-transformed squared coefficient of variation ($CV^2$) of TPM-normalized decomposed tag cluster expression across the LCL panel
- adjusted_log10_CV2: $\log_{10}$-transformed squared coefficient of variation ($CV^2$) after adjustment of the mean expression-dispersion relationship for decomposed tag cluster..
- broad_name: genomic coordinates of the original encompassing promoter provided as chromosome:start-end;strand
- broad_adjusted_log10_CV2: $\log_{10}$-transformed squared coefficient of variation ($CV^2$) after adjustment of the mean expression-dispersion relationship for the original encompassing promoter.
- broad_adjusted_quartile: adjusted variability of the original encompassing promoter split by quartiles (<Q1: 0-25%, (Q1,Q2]: 25-50%, (Q2,Q3]: 50-75%, >Q3: 75-100%)

**Supplementary Table 6: Lead prQTL hit for each promoter.**

[tab-delimited]

Row names: row number

Columns:

- SNP: lead SNP (lowest FDR) genomic coordinates identified as prQTL for the corresponding promoter.
- gene: genomic coordinates of promoter associated with given lead SNP provided as chromosome:start-end;strand
- beta: Effect size estimate for the given SNP-promoter pair
- t.stat: Test statistic (t-statistic) for the given SNP-promoter pair
- p.value: p-value for given SNP-promoter pair
- FDR: False discovery rate estimated using Benjamini–Hochberg procedure

**Supplementary Table 7: Lead frQTL hit for each promoter.**

[tab-delimited]

Row names: row number

Columns:

- broadID: genomic coordinates of original promoters overlapping decomposed promoters associated with frQTL provided as chromosome:start-end;strand
- decomposedID: genomic coordinates of the decomposed promoters with the strongest association (FDR) with given lead SNP provided as chromosome:start-end;strand.
- geneID: Ensembl ID of the associated gene
- SNP: lead SNP (lowest FDR and FDR<=5%) genomic coordinates identified as frQTL for the corresponding promoter.
- beta: Effect size estimate for the given SNP-decomposed promoter pair
- FDR: False discovery rate estimated using Benjamini–Hochberg procedure
- relative_effect: relative change in original promoter expression between major and minor allele for given SNP. Calculated as (A-B)/A using mean TPM-normalized expression for A (homozygous for major allele) and B (homozygous for minor allele).
- adjusted_quartile: adjusted variability of original promoters split by quartiles (<Q1: 0-25%, (Q1,Q2]: 25-50%, (Q2,Q3]: 50-75%, >Q3: 75-100%)

**Supplementary Table 8: Gene level expression characterization across LCLs using RNA-seq (Geuvadis).**

[tab-delimited]

Row names: Ensembl ID of the associated gene

Columns:

- symbol: gene symbol (HGNC ID)
- median: median TPM-normalized gene expression across the LCL panel
- CV: coefficient of variation of TPM-normalized gene expression across the LCL panel
- mean: mean TPM-normalized gene expression across the LCL panel
- log10_CV2: $\log_{10}$-transformed squared coefficient of variation ($CV^2$) of TPM-normalized gene expression across the LCL panel
- adjusted_log10_CV2: $\log_{10}$-transformed squared coefficient of variation ($CV^2$) after adjustment of the mean gene expression-dispersion relationship.

**Supplementary Table 9: Promoters associated with stabilizing frQTLs.**

[tab-delimited]

Row names: row number

Columns:

- broadID: genomic coordinates of original promoters overlapping decomposed promoters associated with stabilizing frQTL provided as chromosome:start-end;strand
- geneID: Ensembl ID of the associated gene
- SNP: Lead SNPs identified as stabilizing frQTLs for given promoter
